# Evolutionary and ontogenetic shifts from aquatic to terrestrial light environments in frogs provide new insights into the vertebrate phototransduction cascade

**DOI:** 10.1101/2025.10.31.685797

**Authors:** Taegan Perez, Rayna C Bell, Camille Lavoie, Matthew K Fujita, David J Gower, Jeffrey W Streicher, Kate N Thomas, Ryan K Schott

## Abstract

Evolutionary studies of vertebrate vision have mainly focused on the light-sensitive opsins neglecting the downstream phototransduction cascade. Here, we investigate patterns of gene loss and expression across jawed vertebrates, providing clarity on the evolution of phototransduction. We next focus on an ecologically diverse sample of frogs represented by whole-eye transcriptomes and genomes from 113 species. We tested the hypothesis that phototransduction genes were lost in the common ancestor of frogs due to nocturnality and that functional differences in phototransduction are driven by variation in habitat, activity pattern, and life history. Across frogs, we recovered 38 of 39 vertebrate phototransduction genes, with all but one of these (*GUCA1B2*) consistently expressed in frog eyes. This contrasts the high level of gene loss found in other ancestrally nocturnal tetrapods (*e.g.*, mammals and snakes) and is more similar to ray-finned fishes. More than other ecological traits, we found that loss of the larval aquatic life stage (direct development) and inhabiting aquatic and semiaquatic habitats as adults were most strongly associated with shifts in selective strength, with widespread signals of both positive and relaxed selection across frog phototransduction genes. These findings suggest that phototransduction is under different functional constraints in aquatic versus terrestrial light environments and that shifts in the use of these environments have played a strong role in frog visual evolution. Collectively, these results reinforce that frogs are of particular interest for vertebrate visual evolution because they span both an evolutionary and an ontogenetic transition from vision underwater to vision on land.

## Introduction

Sensory systems play a critical role in allowing animals to perceive and respond to their environments by detecting and processing external stimuli (Dungan et al. 2016). Among sensory systems, the visual system typically supports a wide range of complex functions that contribute to both survival and reproductive success. The visual system also demonstrates remarkable diversity and specialization across taxa, reflecting adaptations to a wide variety of ecological niches (Nilsson, 2021). By constructing images of the environment, the visual system allows animals to orient themselves within their habitats, locate and identify potential food sources, recognize conspecifics or potential mates, and detect and evade predators. The ability to process visual information near instantaneously provides a significant evolutionary advantage in dynamic and unpredictable environments (Bowmaker, 2008; Cronin et al. 2014).

At the core of this visual capability is the process of photoreception, which begins when a visual pigment (composed of an opsin protein bound to a light sensitive chromophore) is activated by the absorption of a photon of light. Ancestrally, vertebrates express five classes of visual opsins that each form visual pigments maximally sensitive (λ_max_) at different portions of the light spectrum, and whose activation ultimately results in vision. These visual opsins are found in specialized, outer retinal photoreceptor cells in vertebrates and are divided into two types: rods, which are specialized for vision in low-light conditions, and cones, specialized for vision in bright-light conditions. To mediate vision in low-light conditions, rods are highly light sensitive, integrating incoming light over longer periods of time with low noise; however, they have slower activation and recovery rates as a result (Ingram et al. 2016; Lamb, 2016, 2022). Cones are much less sensitive, and thus cannot operate under low light, but have fast response and recovery times (Lamb, 2022). In addition, cones have greater temporal resolution than rods, an adaptive property that is important in brighter and potentially more variable light environments (Ingram et al. 2016). The differences between rod and cone physiology are due, in part, to a unique evolutionary system in which distinct gene sets are employed for the shared task of light detection. The visual opsins present a clear example. Rod photoreceptor cells typically express visual pigments containing rod opsin (*RH1*), maximally sensitive at ∼500 nm whereas the visual pigments found in cone photoreceptor cells typically contain one of the other four visual opsins: long wavelength sensitive (*LWS*; λ_max_ 510–570 nm; green to orange/red light), short wavelength sensitive 1 (*SWS1*; λ_max_ 358– 430 nm; UV to violet/blue light), short wavelength sensitive 2 (*SWS2*; λ_max_ 437–460nm; violet to blue light) or rhodopsin-like 2 (*RH2*; λ_max_ 470–510; blue to green light). The expression of multiple classes of cone opsins in vertebrate retinas is also what enables color vision (Bowmaker, 2008).

In vertebrates, most molecular research on visual system adaptation to photic environments has focused on the early stages of photoreception, namely in the opsin proteins (*e.g.*, Boyette et al. 2024; Macpherson et al. 2024; Schott et al. 2024; Van Nynatten et al. 2024; Castiglione et al. 2023; Schott et al. 2018; Wu et al. 2016). Research has demonstrated various ways in which the vertebrate visual system has adapted to different photic environments through mechanisms such as alterations at key amino acid sites (Van Nynatten et al. 2015; Carlton et al. 2020), gene loss and duplication (Hauser & Chang, 2017; Musilova et al. 2019), and shifts in expression across various taxa and environments (Hauser & Chang, 2017; Schott et al. 2022a).

Much less attention, however, has been paid to the downstream steps of photoreception, notably the remaining steps of the vertebrate phototransduction cascade.

Following light absorption by the visual pigment, the next stage of photoreception in vertebrates is the complex signal transduction cascade (phototransduction) where the light signals are converted to electrical signals. These signals are eventually transmitted through the retina and to the brain for processing of visual information (Box 1; Fig. 1; Larhammar et al. 2009; Palczewki, 2014). Rod and cone photoreceptor cells employ the same core phototransduction cascade across vertebrates but differ markedly in activation and recovery kinetics, and in signal amplification. Rods are specialized for sensitivity, capable of detecting single photons due to a high degree of biochemical amplification: one activated rod opsin can activate multiple transducin molecules, hydrolyzing many cGMP (cyclic guanosine monophosphate) molecules, amplifying visual signals (Arshavsky & Burns, 2012; Lamb & Hunt, 2018). These photon responses are also longer lived in rod cells, due to slower inactivation of the activated opsin and other downstream activation phase proteins (Burns & Pugh, 2010). Cone cells, on the other hand, have markedly lower transducin activation rates and overall lower biochemical amplification (Kefalov, 2012; Kawamura et al. 2017). Cones trade sensitivity for a more rapid response by activating guanylyl cyclase proteins more quickly through elevated expression of guanylyl cyclase activating proteins (GCAPs), enabling faster cGMP synthesis (Lamb & Hunt, 2018). The specialized nature of their open outer segment discs also promote rapid Ca^2+^ dynamics (Kefalov, 2012). Finally, the recovery phase time in cones is also faster than in rods. Cone opsins are phosphorylated more rapidly, and subsequent recovery phase steps are completed rapidly by increased expression of shut-off proteins like those involved in the RGS9 binding complex (Burns & Pugh; Arshavsky & Burns, 2012). Many of these differences in activation and recovery kinetics, and signal amplification, are due, in part, to differences between the cone-and rod-specific isoforms in phototransduction proteins (Fig. 1; Box 1; Schott et al. 2018; Lamb & Hunt, 2018; Lamb, 2020). This system is thought to have emerged from the two rounds of whole genome duplication early in vertebrate evolution (Lamb & Hunt, 2018), providing opportunities for diversification in the functions of the original and/or duplicated copies of the phototransduction genes (Lamb, 2020). Further details on the process of phototransduction and the genes and proteins involved are described in Box 1.

### Box 1. Description of Vertebrate Phototransduction Cascade in Rods and Cones

Here, we describe the vertebrate phototransduction cascade as occurring in the three stages: the dark state, light activation, and recovery.

**I) The Dark State.** This stage describes the state of an inactive photoreceptor. At this time the levels of cytoplasmic cyclic guanosine monophosphate (cGMP) are high. cGMP binds to cyclic nucleotide gated (CNG) channels (rods, CNGCα1 + CNGCβ1; cones, CNGCα3 + CNGCβ3), keeping these channels and Na+/Ca+ exchanger channels (rod, NCKX1; cone, NCKX2) open and allowing the flow of Na^+^ and Ca^2+^ ions into the photoreceptor cell. This depolarization causes a continuous release of the neurotransmitter glutamate at the synaptic terminal, an inhibitory signal to downstream neurons.
**II) Light Activation.** This stage begins the process of converting a light signal into an electrical signal. This begins with the absorption of a photon of light by the retinal chromophore component of the visual pigment, triggering a *cis* to *trans* isomerization. This photoisomerization causes a conformational change in the opsin protein (rod: RH1; cones, LWS, RH2, SWS1, SWS2) of the visual pigment to its active state. The activated visual pigment subsequently activates a heterotrimeric G protein called transducin. The binding of Mg^2+^-GTP and release of GDP causes the dissociation of transducin into its individual alpha (rod, Gαt1; cone: Gαt2), beta (rod, Gβ1; cone, Gβ3), and gamma (rod, Gγt1; cone, Gγt2) subunits. The alpha-subunit of transducin further relays this signal of light activation by stimulating the activity of a cGMP-phosphodiesterase, PDE6 (rod, PDEα, PDEβ, PDEγ; cone, PDEα’, PDEγ’). During the dark state, PDE6 is kept inactive by the γ-subunits, but binding of activated alpha transducin subunits remove this inhibition. The activation of PDE6 results in a decrease in the concentration of cGMP and intradiscal Ca^2+^ leading to a closure of the CNG channels, hyperpolarizing the cell. Closure of CNG channels reduces the influx of Na^+^ and Ca^2+^ ions, decreasing cytosolic Ca^2+^ as well. Hyperpolarization of the cell reduces glutamate release at the synapse, allowing the cells’ generated electrical response to travel through the retina to the brain (Lamb, 2020).
**III) Recovery.** The reduction and termination of signaling in the phototransduction cascade is first mediated by the phosphorylation of the opsin protein via a G-protein-coupled receptor kinase, GRK (rod, GRK1A; cone, GRK1B, GRK7). GRK activity is regulated by recoverin (rod, Rec; cone, Visinin), a Ca^2+^ binding protein that inhibits GRK in darkness. When light closes CNG channels, causing a drop in cytosolic Ca^2+^ concentration, recoverin releases GRK, and the kinase becomes active, binding to the opsin and reducing further transducin activation. Further G-protein activation is stopped by subsequent binding of arrestin (rod: Arr-S; Cone, Arr-C). Meanwhile any activity of transducin is shut off by GTP hydrolysis to GDP, using Mg^2+^ bound to the nucleotide. Binding of the regulator of G protein signaling 9 complex that includes RGS9, a RGS9 binding protein (R9AP, R9AP-B or R9AP-like, and G protein β (Gβ5) accelerates this process. The inactivation of transducin also inactivates PDE6 as the inhibitory subunits reengage the catalytic subunits. cGMP regeneration begins as guanylyl cyclase (GC) stimulated by GC-activating proteins, GCAPs (both: GCAP1, GCAP1L, GCAP2, GCAP2L; cone, GCAP3). GC catalytic activity is Mg^2+^ dependent, with Mg^2+^-GTP serving as the substrate. GCAPs are Ca^2+^ sensing proteins, as Ca^2+^ concentration falls, GCAPs switch to an activating confirmation, increasing GC activity. The newly synthesized cGMP reopens CNG channels, allowing Na^+^ and Ca^2+^ influx, depolarizing the photoreceptor cell, increasing glutamate release, and returning the cell to the dark state. The opsin component of the visual pigment is also returned to their ready state by a return of the all-*trans*-retinal to an 11-*cis*-retinal through the visual cycle (Larhammar et al. 2009).

**Fig. 1.**
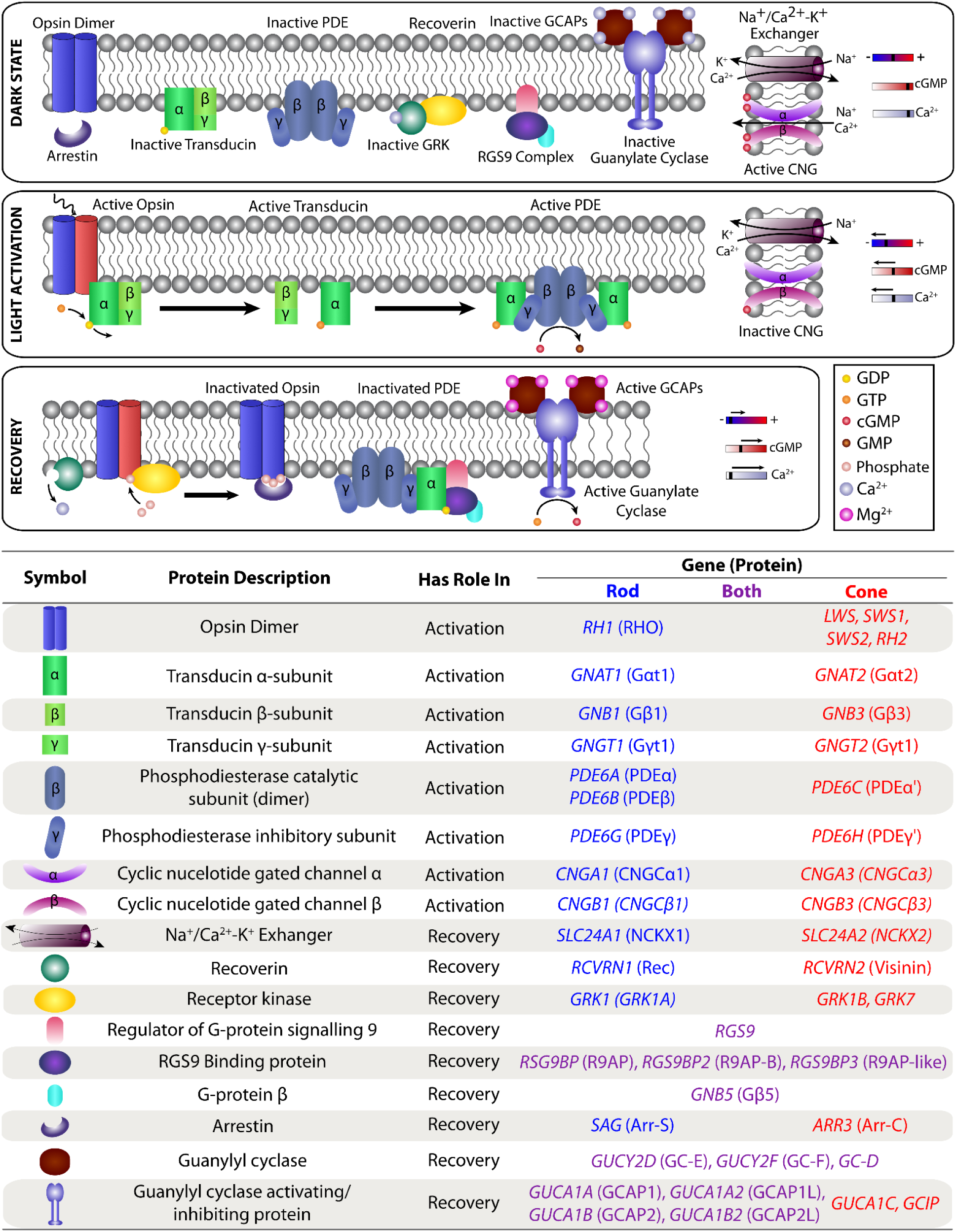
Typical Vertebrate Phototransduction Cascade. The Dark State refers to the inactivated state of the photoreceptor cell. Following the absorption of a photon of light by the visual pigment (opsin bound to a vitamin A chromophore), the photoreceptor enters the light activation phase. During the recovery phase, the opsin is returned to its inactive state. The photoreceptor returns to the dark state when the retinal based chromophore is regenerated through the visual cycle.

End of box

Despite the importance of the phototransduction cascade in transmitting and amplifying light signals, the striking physiological difference between rods and cones, and the adaptive potential of rod-and cone-specific phototransduction genes, relatively few studies have examined vertebrate phototransduction gene evolution. However, some studies have found limited evidence for adaptation in particular vertebrate lineages. For instance, positive selection in mammalian rod-specific phototransduction genes has been linked to an inferred shift to nocturnality in ancestral mammals (Wu et al. 2017). In snakes, evidence for positive selection was disproportionately found in cone-specific phototransduction genes, which was proposed to result from repeated shifts from nocturnal to diurnal activity patterns (Schott et al. 2018). Likewise, evidence for positive selection among rod-specific phototransduction genes in owls is thought to have resulted from the shift to a nocturnal activity pattern from the diurnal pattern exhibited by most other birds (Wu et al. 2016).

Collectively, these examples suggest that shifts in activity patterns may be a driving force in tetrapod phototransduction gene evolution, but studies are lacking from other vertebrate lineages, including those that use aquatic habitats, such as fishes and amphibians.

Amphibians are of particular interest for visual evolution because they span both evolutionary and ontogenetic transitions from vision underwater to vision on land. Frogs and toads (Order: Anura, hereafter collectively “frogs”) represent the most diverse amphibian group with over 7800 extant species described to date, representing more than 200 million years of adaptation to both terrestrial and aquatic environments (AmphibiaWeb, 2025). The visual system is critical to many aspects of frog biology including foraging, movement patterns, and habitat preferences (Buchanan 2006), and correspondingly, elements of both life history and light environment have influenced the evolution of frog visual systems. For instance, frogs are inferred to be ancestrally nocturnal, and while most frog species maintain a nocturnal activity pattern, several lineages have independently evolved diurnality (Anderson & Wiens, 2017). These repeated transitions to diurnality are associated with differences in morphology (Thomas et al. 2020, Shrimpton et al. 2021, Mitra et al. 2022) and optics of the visual system (Yovanovich et al. 2019, Yovanovich et al. 2020, Thomas et al. 2022). Although evolutionary transitions between diurnal and nocturnal activity patterns are associated with selective shifts and positive selection in vision genes for other tetrapod groups (Wu et al. 2016, 2017; Schott et al. 2018, 2019; White et al. 2022), thus far there is limited evidence for a relationship between transitions to diurnal activity and shifts in selection for visual opsins (Schott et al. 2022b, Wan et al. 2023, Schott et al. 2024) or non-visual opsins in frogs (Boyette et al. 2024).

Most frogs have a biphasic life cycle with an aquatic larval stage that metamorphoses into a terrestrial adult, which requires the visual system to operate in two very different light environments within the lifespan of a single individual. Several independent lineages of frogs, however, have lost the tadpole stage and instead hatch directly from eggs with miniature adult morphology (*i.e.*, as froglets). Both visual opsins (Schott et al. 2024) and non-visual opsins (Boyette et al. 2024) exhibit shifts in selective pressure for frogs with direct development relative to those with aquatic larvae, with evidence that these shifts are due to a relaxation of selective constraint.

Because the direct-developing species lack an aquatic tadpole stage, relaxation of selective constraint may reflect that they are not constrained to a dual functional role of their visual system in both water and air. Whether relaxation of selective constraint in direct-developing frogs extends beyond visual opsin genes has yet to be tested. Similarly, the ecological diversity of frogs is associated with differences in both morphological and genomic aspects of the visual system. For instance, fossorial and aquatic species have smaller eyes, whereas scansorial (climbing) species have greater eye investment and lower UV light transmission through the lens, which is proposed to benefit high temporal resolution and acuity while navigating complex three-dimensional environments (Thomas et al. 2020, 2022). Furthermore, scansorial frogs exhibit differences in selective pressure in the visual opsin *RH1* that may result in faster temporal resolution for scotopic vision and be beneficial in navigating complex arboreal environments (Schott et al. 2024).

Here we aim to characterize phototransduction gene evolution across the frog tree of life using a dataset extracted from *de novo* assembled frog eye transcriptomes (Schott et al. 2022, 2024), and from available genome assemblies, that together represent 113 species and 34 of the 57 currently recognized frog families (AmphibiaWeb, 2025). Because research on the evolutionary history of phototransduction genes and proteins, both across and within vertebrate lineages, remains limited, we first aim to identify the broad patterns of gene retention and loss across major jawed vertebrate lineages using the recent increase in available genomic resources, and assess their expression in the eye through transcriptomic analyses. We expect that several genes involved in the vertebrate phototransduction cascade previously thought to be absent based on more limited data are not only present in major vertebrate lineages, including frogs, but are also actively expressed in the eye. Within this broad context, we test the hypothesis that a subset of phototransduction genes were lost in the common ancestor of extant frogs as demonstrated in other vertebrate groups with nocturnal ancestry (Castoe et al. 2013; Schott et al. 2018, 2019; Wu et al. 2017; Gemmel et al. 2020).

Finally, we investigate the molecular evolution of frog phototransduction genes using codon-based models to test for signatures of positive selection and for long-term shifts in selective pressure associated with variation in light availability and quality due to differences in activity patterns, life history, and habitats. Based on studies in other tetrapods with diurnal and nocturnal transitions (Wu et al. 2016, 2017; Schott et al. 2018, 2019; White et al. 2022), we might expect to find patterns of positive selection in diurnal frog species and shifts in selection pressure between diurnal and nocturnal species, especially in cone-specific genes. Previous studies of molecular evolution of visual opsins and non-visual opsins in frogs, however, have not found activity pattern to be a strong driver of selective pressure and instead found that differences in life history and habitat were linked most strongly to the evolution of these genes (Boyette et al. 2024; Schott et al. 2022, 2024; but see Wan et al. 2023). For instance, aquatic and fossorial frogs have smaller eyes, and their cone-specific visual opsins are under relaxed selective constraints with hints of potential adaptive evolution (Thomas et al. 2020; Schott et al. 2024). By contrast, scansorial frogs have larger eyes with selective shifts in *RH1* sequences that could indicate selection for increased temporal resolution through faster RH1 kinetics (Thomas et al. 2020; Schott et al. 2024). Based on these prior findings, we predict that scansorial frogs will show signals of adaptive evolution broadly across their phototransduction genes if coevolutionary changes had occurred to enhance visual acuity, or these signals would be concentrated in rod-specific genes if changes to temporal resolution had occurred at the level phototransduction gene sequences. By contrast, if fossorial and aquatic species have divested in the visual system, we predict a relaxation of selective constraint on their phototransduction genes with more relaxation in cone-specific genes. Additionally, we test the importance of species distribution in the phototransduction gene evolution of frogs. Tropical species are typically exposed to less variation in photoperiod than their temperate counterparts (Borah et al. 2019). The variation in available light has implications for the evolution of the light signaling cascade. Specifically, we expect to see tropical species experience a relaxation of selective constraint, especially in cone-specific phototransduction genes because of the more consistent photoperiod. Finally, we test the hypothesis that the loss of the aquatic tadpole stage in direct-developing frogs resulted in a release of selective constraint for phototransduction to function in both aquatic and terrestrial light environments.

## Results

### Recovery and Expression of Jawed Vertebrate Phototransduction Genes

We recorded the presence and absence of genes associated with vertebrate phototransduction in the genomes of all major jawed vertebrate lineages with a focus on tetrapods (Fig. 2, Supplementary File S1). This included 39 known and putative vertebrate non-opsin phototransduction genes and the five vertebrate visual opsins. Three genes were flagged for more detailed analysis due to their unconfirmed roles in phototransduction and inconsistencies with recoveries previously reported in the literature. These included the “olfactory” guanylyl cyclase (*GC-D*), inhibitory guanylyl cyclase protein (*GCIP*), and a putative isoform of the inhibitory subunit of phosphodiesterase, *PDE6I*. We also investigated expression of the guanylyl cyclase activity protein isoform *GUCA1B2* among jawed vertebrates given the lack of expression of this gene in frog eyes.

**Fig. 2.**
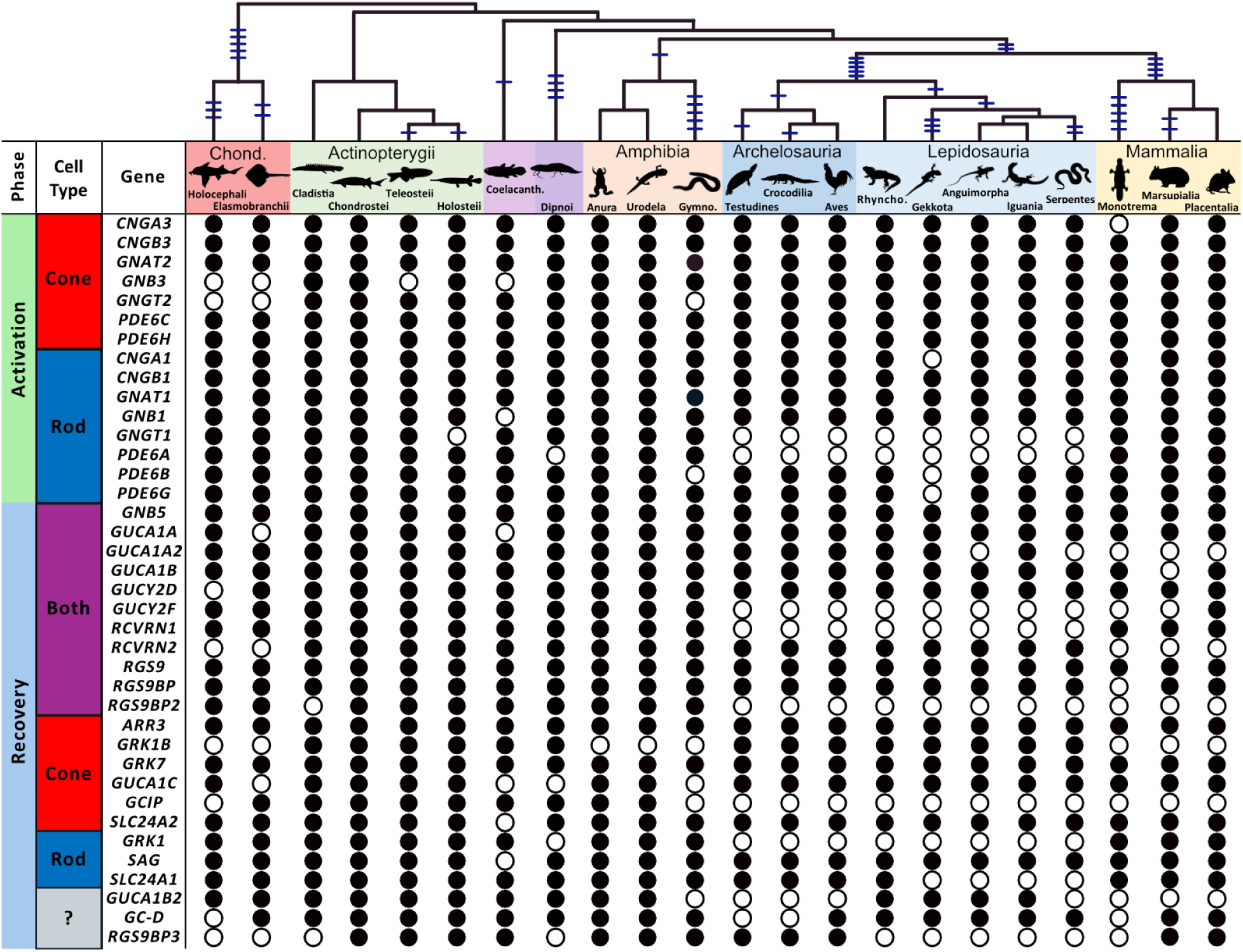
Presence and absence of phototransduction genes in major jawed vertebrate lineages. Evidence for the presence of a gene in a descendant lineage (*i.e*., present in at least one member) is shown with a filled circle while absence from all tested species is shown with an open circle. The phylogenetic relationships between vertebrate lineages are shown at the top following Irisarri et al. (2017). Tick marks are shown along branches to indicate the number and inferred location of phototransduction gene losses. The photoreceptor cell type (if known) and phase (activation vs recovery) are also shown. A full list of publicly available genomes and transcriptomes used for the analyses is available in Supplementary File S1. Abbreviations (from left to right): Chond.: Chondrichthyes, Coelacanth: Coelacanthiformes, Gymno.: Gymnophiona, Rhyncho.: Rhynchocephalia.

Overall, we found that major vertebrate lineages are inferred to have specifically lost 1.6 phototransduction genes on average (excluding visual opsins), ranging from 0–5 genes (*i.e.*, genes lost along the branch leading to that lineage; Fig. 2) with total accumulated losses (*i.e.*, specific losses, plus those lost ancestrally) ranging from 0–12 genes (5.6 on average). Lineages inferred as having nocturnal ancestry (Anderson & Wiens, 2017) have lost more phototransduction genes than those with diurnal ancestry, including mammals (4 lineage-specific, 6 in total), snakes (2 lineage-specific, 11 in total), and geckos (3 lineage-specific, 12 in total). Notable exceptions to this pattern are the tuatara (0 specific, 8 total), as previously noted (Gemmel et al. 2020) and frogs and salamanders (0 specific losses, 1 in total), comparable to the losses inferred in lineages of ray-finned fishes (0–2 specific with no ancestral losses).

The “olfactory” guanylyl cyclase (*GC-D*) gene and protein have previously been suggested to be involved in phototransduction (Lamb, 2020). We found complete coding sequences for *GC-D* in most major vertebrate lineages with the exception of turtles, crocodilians, and monotremes. Although the presence of *GC-D* in vertebrate genomes has support, less is known regarding its possible function in vertebrate vision. We checked for expression of *GC-D* in publicly available whole-eye transcriptomes for most major vertebrate lineages and although *GC-D* may be maintained in many vertebrate genomes it is not consistently expressed in vertebrate eyes. We found complete *GC-D* coding sequences in Chondrichthyes, Holostei and Anura (frogs), but only partial sequences in Teleostei. Among reptilian eye transcriptomes, we found complete coding sequences for Aves, but only partial coding sequences for all squamates available for testing.

Finally, despite evidence for the presence of *GC-D* in mammals, we did not find evidence for the expression of *GC-D* in the eye. It appears that *GC-D* is retained across most major vertebrate lineages, but consistent expression in the eye is restricted to frogs and fishes (at least Holostei).

*GCIP* was recovered only in amphibians (frogs and salamanders, but not caecilians), cartilaginous fishes, ray-finned fishes, and lobe-finned fishes (excluding tetrapods) genomes and appears to be lost in all other vertebrate lineages (reptiles and mammals). In terms of eye expression, we unsurprisingly did not find evidence of *GCIP* expression in any reptiles (including birds) or mammals. Among fishes, we found evidence for partial coding sequences of *GCIP* only in teleost fishes but did not find evidence for expression in Chondrichthyes or Holostei. We were unable to test coelacanth or lungfish eye expression due to a lack of available transcriptomic data. Thus, it appears that *GCIP* is maintained in multiple vertebrate lineages, but eye expression is restricted to frogs.

Authors have previously suggested the presence of a third *PDE6* inhibitory subunit gene in vertebrates, *PDE6I*, having originated in the second round of whole genome duplication (Lagman et al. 2016; Lamb, 2020). Lagman et al. (2016) found evidence for *PDE6I* in Teleostei, Holostei (gar), Coelacanthimorpha, and Anura, but note that *PDE6I* was subsequently lost in amniotes. Using these previously identified sequences as queries, we searched for sequences of putative *PDE6I* genes from predicted gene sequences in the NCBI nucleotide database and for expression in our frog and vertebrate-wide eye transcriptome datasets. We found putative *PDE6I* sequences in the genomes of the same four groups as identified previously, and also Cladistia (bichir), Chondrostei (sturgeon and paddlefish), and Dipnoi (lungfish), but not in any other lineages (Fig. S1, Supplementary File S1). Complete and partial sequences were also recovered from 12 of the frog eye transcriptomes (11 species) with expression in both the adult and tadpole of one species (*Ascaphus truei*). Additionally, complete sequences were recovered from one of the eye transcriptomes from vertebrate-wide dataset of another holost fish (*Amia calva*) and one teleost fish (*Astyanax mexicanus*), but not the other two teleosts we tested (*Oreochromis niloticus* and *Astatotilapia burtoninone*) or any of other eye transcriptomes (Supplementary Files S1 and S2). The recovered putative *PDE6I* genes were highly variable in length and their 5’ sequences had very little to no homology after codon alignment, with the 3’ end being highly conserved (Perez at al. 2025).

Phylogenetic analysis of the conserved regions of *PDE6G*, *PDE6H*, and *PDE6I* found that most putative *PDE6I* genes formed a clade sister to *PDE6H*, except that this clade also contained cartilaginous fish *PDE6H*. A second clade of putative *PDE6I* sequences, which contained sequences from sturgeon and paddlefish (Chondrostei) formed a clade sister to the *PDE6H* and other *PDE6I* sequences (Fig. S1). We do not consider the placement of the clades to be reliable due to the short length of the gene. Indeed, during identification of possible PDE6I sequence we recovered several other topologies using similar alignments, including placement of cartilaginous fish *PDE6H* with other *PDE6H* (Taegan et al. 2025). However, the result does support the present of an additional gene, or genes, beyond *PDE6G* and *PDE6H*. Due to its uncertain role in phototransuction, low recovery rates from the eye transcriptomes, and low 5’ homology across taxa, we did not consider *PDE6I* in our analyses of gene loss or molecular evolution.

Due to the lack of expression of the guanylyl cyclase activating protein isoform *GUCA1B2* in frog eyes, we examined the presence and expression of this gene among other jawed vertebrate lineages. We recovered complete coding sequences from the genomes of some vertebrate lineages apart from caecilians, archelosaurs (turtles, crocodiles, and birds), snakes, and mammals. In terms of expression, we recovered partial coding sequences for Chondrichthyes and Teleostei, and complete coding sequences in one species of Holostei and one species of Squamata.

### Phototransduction Gene Recovery in Frogs

Sampling included a total of 113 species, with 83 whole eye transcriptomes and 30 genomes (Fig. 3). A total of 3697 whole or partial coding sequences were recovered representing all 37 phototransduction genes previously documented in frogs. Complete or partial coding sequences were consistently recovered for more than 80% of species across nearly all phototransduction genes (Supplementary File S2). The genes *GUCA1C* and *GUCA1B2* are notable exceptions.

**Figure 3.**
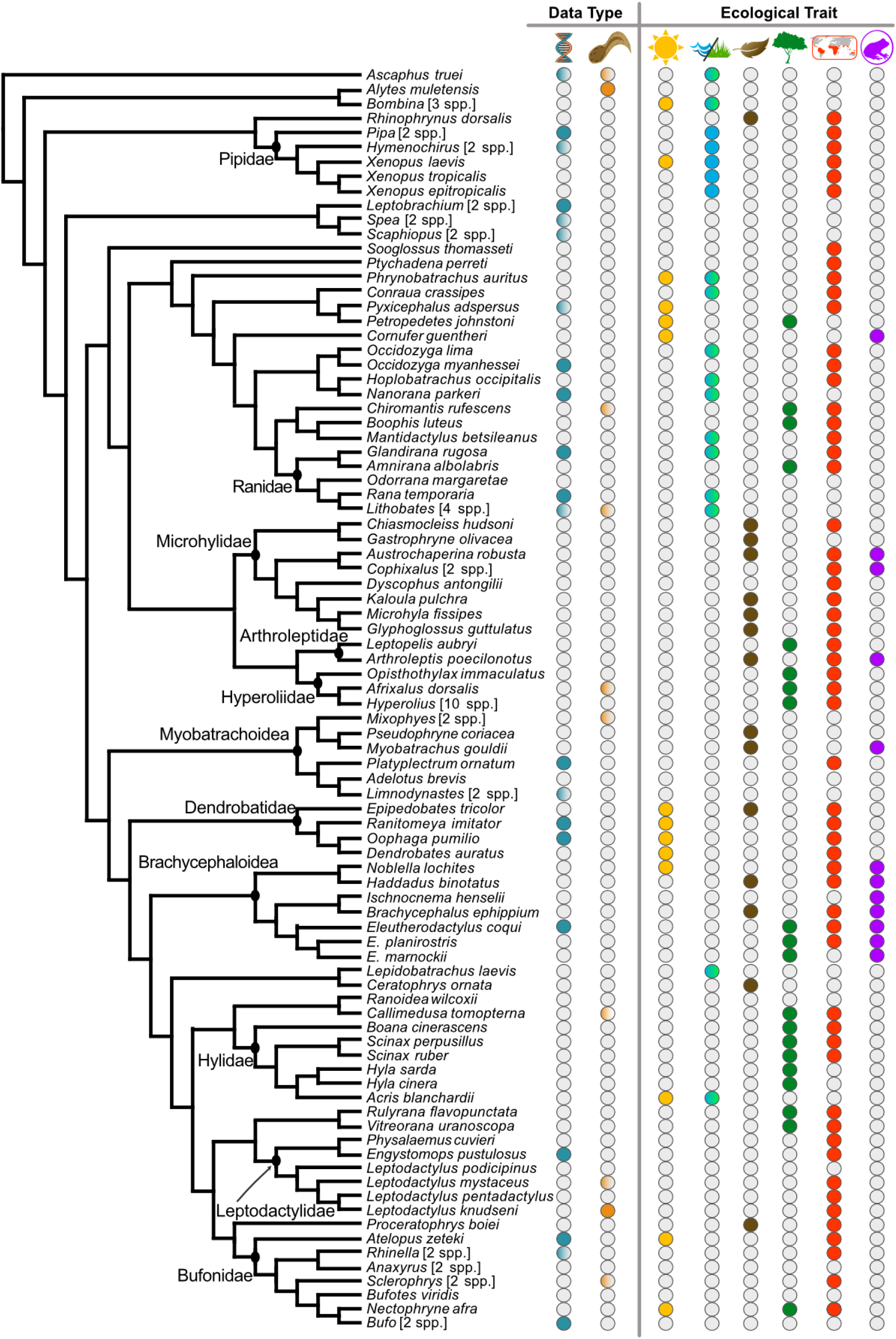
Phylogenetic relationships, data types, and ecological traits of frog species included in the phototransduction gene datasets. Data type classification from left to right (with associated symbols in parentheses): genomic or transcriptomic (DNA helix) and life cycle stage (tadpole). Species represented by genomic data are denoted by a filled circle, and those with both transcriptomic and genomic data are partially filled. Species represented by a tadpole specimen are denoted by a filled circle, and species with both tadpole and adult specimens are partially filled. Unfilled circles indicate transcriptomic data and adult specimens, respectively. Trait classification from left to right (with associated symbols in parentheses) are as follows: adult diurnal activity (sun), adult aquatic or semi-aquatic habit (waves/grass), adult secretive or fossorial habit (leaf), adult scansoriality (tree), tropical distribution (map), and direct development (frog egg). Fully aquatic species are denoted with a solid (blue) circle whereas semi-aquatic species are split (blue/green). Species within the same genus that share the same trait classifications have been collapsed to a single branch for display purposes, and the total numbers of species sampled in those lineages are shown in square brackets. Evolutionary relationships are based upon the phylogenetic hypothesis of Jetz & Pyron (2018). See Schott et al. (2024) for full species sampling information and ecological trait classification.

*GUCA1C* had a recovery rate of 82% in the genomes and 71% in the transcriptomes, and *GUCA1B2* had a recovery rate of 67% in the genomes and 18% in the transcriptomes. For these two genes, no clear phylogenetic pattern of loss was noted. *GUCA1B2* was dropped from downstream analyses because low recovery rates of genes limit success in generating reliable phylogenies and in the performance of the selection analyses. In addition, the lack of consistent expression in the eye transcriptomes suggests this gene may not be involved in visual phototransduction in frogs. The gene *PDE6G* was also dropped from downstream clade model analyses because it is both short (264 bp) and was highly conserved among sampled species. Such genes typically have low phylogenetic signal and are often unable to infer reliable phylogenies as a result (Lamb et al. 2016). Furthermore, models such as PAML that utilize ML methods are subject to greater error when even a few sequences are identical (Yang et al. 2005).

### Evidence for positive selection in a subset of frog phototransduction genes

We found that selection results from both the species tree and gene tree were in concordance and did not change our interpretations, indicating robustness to minor differences in topology. We present results based on our species-tree analyses here, but full results for gene trees are available online (Supplementary File S3). To determine the overall selective constraint acting on frog phototransduction genes, we used the PAML M0 model to estimate the average rate ratio of nonsynonymous to synonymous substitutions (*⍵* or *d*_N_/*d*_S_) across all the codon sites in a given gene alignment. Overall, frog phototransduction genes are under negative purifying selection, with *ω* between 0.003 and 0.195 (Fig. 4). This result was expected given the presumed functional importance of these genes in the frog visual system. The selective constraint in these genes is also in line with values for other functionally important visual genes in frogs, including visual opsins (*ω* = 0.096–0.113; Schott et al. 2024) and non-visual opsins (*ω* = 0.09–0.25; Boyette et al. 2024). We noted no significant patterns in overall selective constraint between either photoreceptor type (*e.g.*, rod vs. cones) or between activation and recovery (Supplementary File S3; Mann Whitney U test, all p>0.05).

**Fig. 4.**
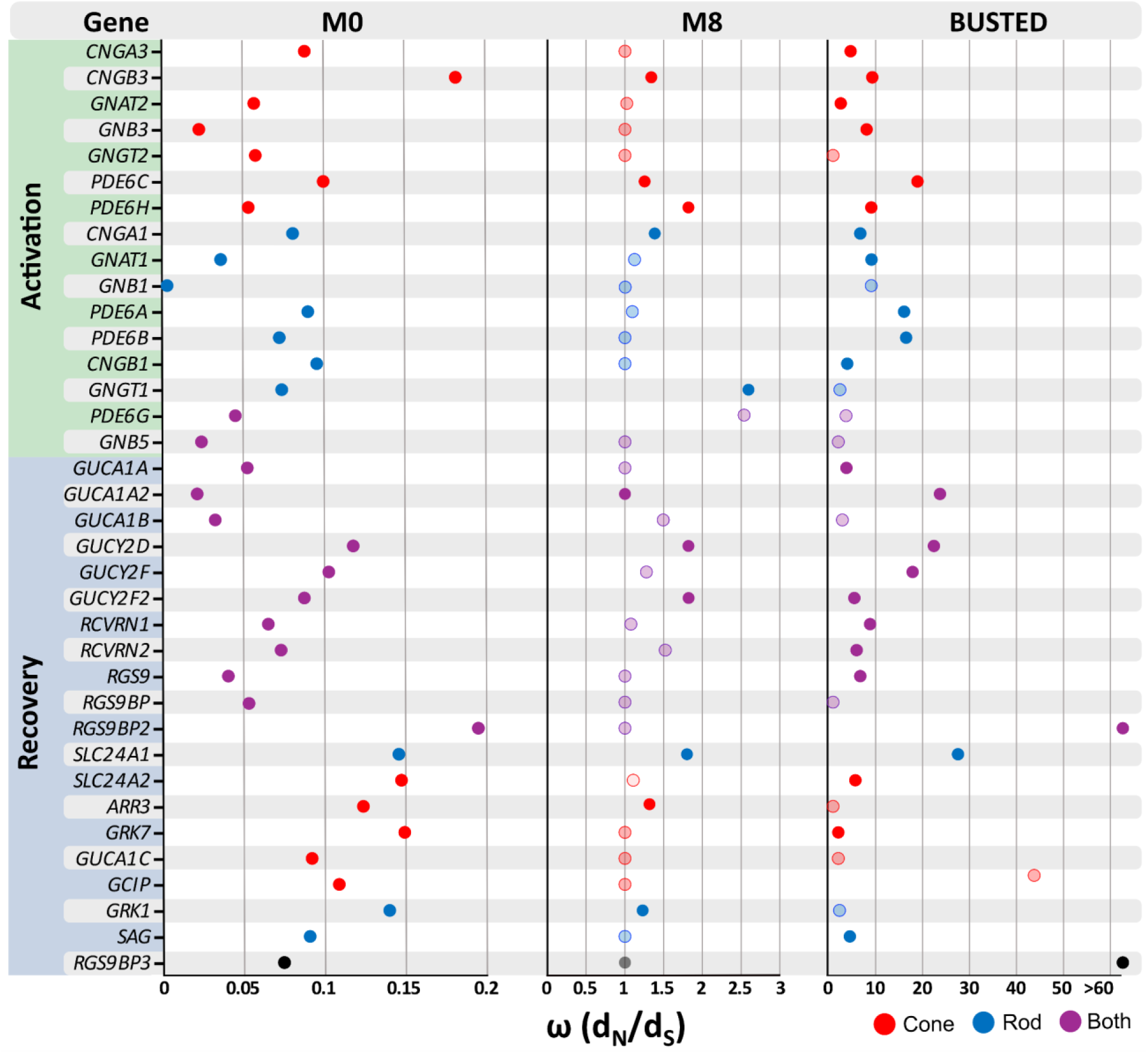
Patterns of selective constraint across frog phototransduction genes. M0: PAML M0 analyses average *ω* across all codon sites in each gene alignment. Within the genes analyzed *CNGB3* and *RGS9BP2* showed the most elevated *ω* values and *GNB1* showed the lowest ω value. The results demonstrate variation in selective constraint across frog phototransduction genes. All *ω* are below 1 indicating overall purifying selection. **M8:** PAML M8 tests for the presence of positively selected codon sites in a gene alignment. The *ω* estimate for the positively selected site class is shown. Eleven genes demonstrated statistically significant evidence for positively selected sites after FDR correction (q<0.05) and are shown with opaque circles. **BUSTED:** HYPHY BUSTED with synonymous rate variation tests for the presence of gene-wide episodic diversifying selection. The *ω* of the third site class of the unconstrained model is shown. Twenty-five genes demonstrate statistically significant evidence for episodic diversifying selection after FDR correction (q<0.05) and are shown with opaque circles.

Genes that demonstrate overall selective constraint may still contain positively selected sites. We tested for this using the PAML M8 model, which allows for *ω* to vary between sites in a gene, and which can be compared to the M7 model (*ω* can vary but is constrained to ≤ 1) using a likelihood ratio test to check for evidence of positively selected sites. Using this method, we found evidence for positive selection in a subset of sites in 11 frog phototransduction genes: four that are typically found only in cones (*CNGB3*, *GRK7*, *PDE6C* and *PDE6H*), three typically found only in rods (*CNGA1*, *GNGT1*, and *GRK1*), and four found in both photoreceptor types (*GUCA1A2*, *GUCY2D*, *GC-D*, and *SLC24A1*) (Fig. 4). A criticism of evidence for positive selection using the PAML M8 model is that this model is not able to account for synonymous rate variation across a phylogeny (Wisotsky et al. 2020). Because PAML M8 estimates *ω* as a single parameter, *ω* can be overestimated when synonymous rate varies across a phylogeny. To address this, we also tested our alignments using the BUSTED model (Murrell et al. 2015) on the Datamonkey HyPhy server (Delport et al. 2010), which allows for synonymous rate variation. We found 25 genes with statistically significant evidence of episodic diversification (positive selection; p<0.05; Fig. 4; Supplementary File S4). This included 9 of the 11 genes identified using the M8 model in PAML. *GRK1* and *GNGT1* were found to be significant using PAML M8 but not BUSTED with synonymous rate variation. Both *GRK1* and *GNGT1* were found to have significant evidence of episodic diversification with BUSTED when synonymous rate variation was not accounted for. This suggests that the evidence for positive selection seen in these genes under the PAML M8 model may be driven by synonymous rate variation rather than nonsynonymous rate variation. Previous studies have attributed the differences between BUSTED and PAML M8 to be primarily related to differences in the model formation between the two methods rather than being evidence for false inferences of positive selection (Boyette et al. 2024). Given the differences in results between PAML and BUSTED, this highlights the importance of multiple model support for selection analyses. After false discovery rate (FDR) correction, all PAML and BUSTED test results remained significant (Supplementary File S3). Complete PAML and BUSTED results can be found in supplementary table S4 and S5, Supplementary Materials online.

### Shifts in Selective Constraint among Phototransduction Genes are Associated with Variation in Frog Ecological Traits

Tests for shifts in selective constraint acting on frog phototransduction genes were completed using PAML CmC models with different ecological partitions (activity pattern, habit, geographic distribution, and life history; Fig. 3). For each gene, we found a significant difference in selection pressure with at least one of the partitions tested (Fig. 5), and except for *GNGT2*, the best-fitting partition was the same using either the species-or gene-tree topology (Supplementary File S4). Significant differences in selective constraint were found for many of the trait partitions (Fig. 5). The direct-developing partition was the best-fitting partition for the greatest number of genes (*CNGA1*, *CNGA3*, *CNGB1*, *GRK7*, *GUCA1A2*, *GUCY2F*, *GC-D, PDE6A, PDE6B,* and *SAG*). This indicates that within each of these genes, the difference in selective constraint was best explained by the direct-developing partition, compared to the other partitions that were tested. In the case of the direct-developing partition, direct-developing species had a greater *ω* compared to non-direct developing species. The fully + semi-aquatic partition was the best-fitting partition for the second-greatest number of genes (*ARR3, GNB3, GNGT1, GUCA1B, GUCY2D, PDE6C, PDE6H, RGS9BP,* and *SLC24A2*), followed by fully aquatic for eight genes (*GNGT2, GRK1, GUCA1A, GUCA1C, RCVRN1, RCVRN2, RGS9,* and *SLC24A1*). The remaining traits were the best fit for a small number of genes: tropical for three genes (*CNGB3*, *GCIP*, *GNB5*), scansorial for two genes (*GNAT2*, *RGS9BP3*), fossorial for one gene (*RGS9BP2)* (Fig. 5). Surprisingly, the diurnal partition was significant for ten genes but was not the best fit for any. Selection analyses were robust to slight differences in topology between ML gene trees versus species trees apart from *GNGT2,* where under the gene-tree topology the fully + semi aquatic partition was the best fit and under the species-tree topology the fully aquatic partition was the best fit. FDR corrections showed evidence for possible false positives in three previously significant results, but none of these was the best-fitting partition for that particular gene (Supplementary File S3).

**Fig. 5.**
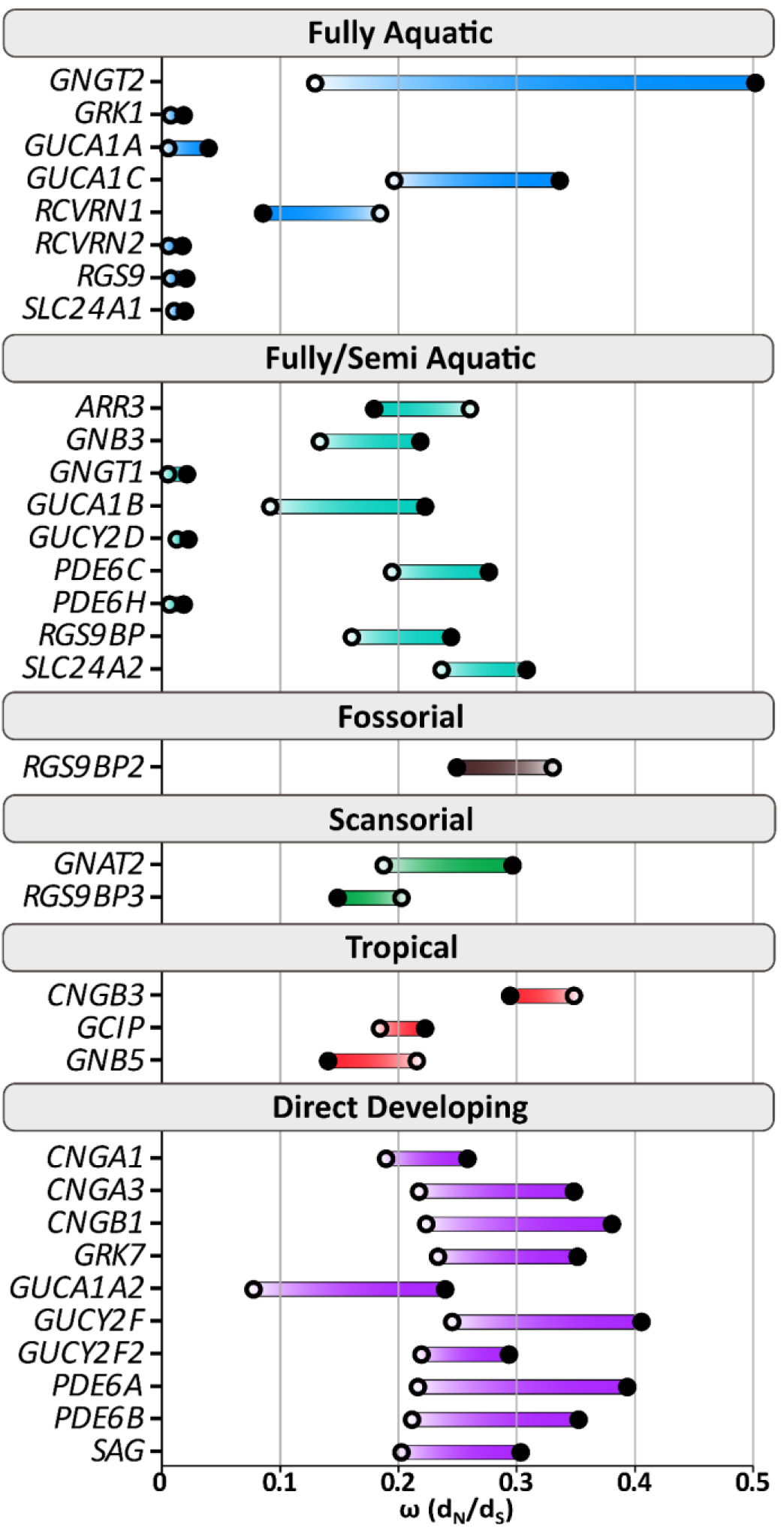
Shifts in selective pressure on frog phototransduction genes across partitions illustrated in. **Figure 3**. Only the best-fitting partition for each of the genes is shown. The *ω* (*d*_N_/*d*_S_) values of the divergent site class using CmC analysis of phototransduction genes with the species-tree topology are shown and highlight the difference between the background *ω* (unfilled circles) and the foreground *ω* (filled circles). Complete PAML CmC results are available in Supplementary File S4.

To determine whether the shifts in selective pressure identified using CmC are due to relaxed selective constraint or potential adaptative evolution we used the RELAX and BUSTED models (Table 1). With the direct-developing partition, the RELAX analyses demonstrated significant evidence for relaxed selection acting on *CNGB1*, *GRK7*, *GUCY2F*, *GC-D*, *PDE6A*, *PDE6B*, and *SAG*; however, only *CNGB1* and *GC-D* showed evidence for positive selection with BUSTED (Table 1, Supplementary Files S6 and S7). The direct-developing partition also showed significant evidence for intensification of selection in *GUCA1A2*, but not positive selection with BUSTED (Table 1, Supplementary Files S6 and S7). Within the fully aquatic and fully + semi-aquatic partitions there was significant evidence for relaxation of selection in *GNB3*, *GUCA1A*, *GUCA1B*, *PDE6C*, *RGS9*, and *RGS9BP*; however, only *GNB3*, *PDE6C*, and RGS9 showed evidence for positive selection with BUSTED (Table 1, Supplementary Files S6 and S7). The fully aquatic and fully + semi aquatic partitions also showed evidence for the intensification of selection in *PDE6H*, *RCVRN1*, and *SLC24A1*; and *PDE6H* and *SLC24A1* also showed evidence for positive selection with BUSTED (Table 1, Supplementary Files S6 and S7). The tropical partition showed evidence for the intensification of selection in *GNB5*; and evidence for positive selection in *CNGB3* with BUSTED (Table 1, Supplementary Files S6 and S7).

**Table 1.**
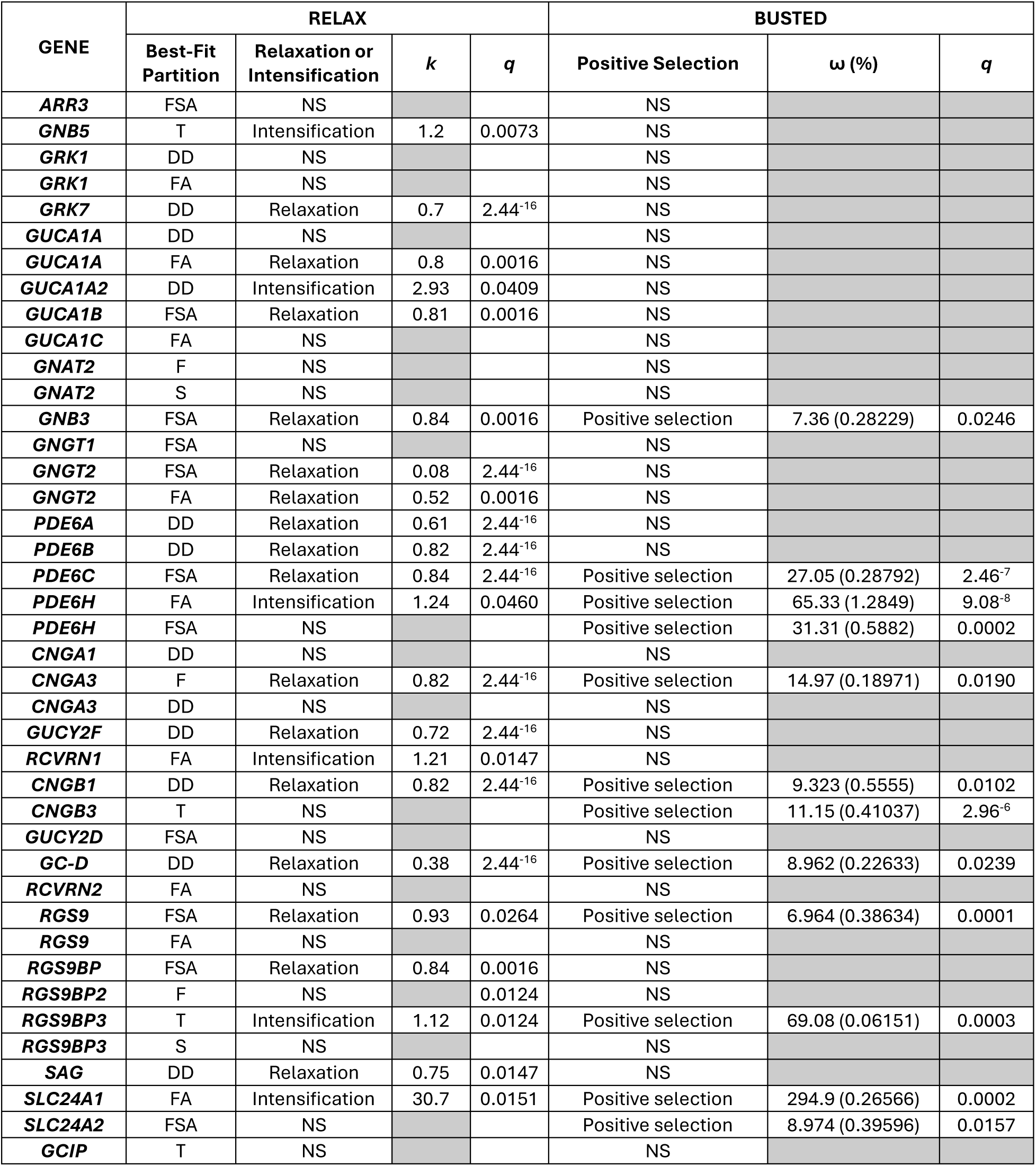
RELAX and BUSTED model results for the best-fitting partitions from the CmC analyses. Abbreviations: FA, fully aquatic; **FSA**, fully and semi-aquatic; **F**, fossorial; **NS**, not significant; **S**, scansorial; **T**, tropical; **D**, direct developing. Full RELAX and BUSTED model results can be found in Supplementary Files S6 and S7.

## Discussion

We investigated the presence and expression of phototransduction genes and proteins across vertebrate lineages with a particular focus on frogs, which have remained relatively understudied in terms of the evolution of genes involved downstream in the phototransduction cascade. To determine which genes are present and expressed in frogs and other vertebrates we used a combination of *de novo* assembled transcriptomes from whole frog eyes and available genomic and transcriptomic resources for frogs and other vertebrates. We obtained phototransduction gene sequences from 113 frog species across 34 families and recovered 38 out of 39 genes with only two genes having low recovery (*GUCA1B2* and *PDE6I*). This result indicates that frogs retain one of the most complete phototransduction repertoires among vertebrates despite the inferred nocturnal ancestry of the group. Across frogs, we uncovered patterns of positive selection in a large subset of the phototransduction genes, with strong evidence for variation in selective constraint between lineages partitioned across different ecologies, behaviors, and life-history strategies. We found that shifts between (semi-)aquatic and other habits, and between direct-developing and biphasic life histories, best explained the variation in selective constraint across species, possibly indicating functional adaptation in frog phototransduction genes. We discuss these findings below in conjunction with our current understanding of phototransduction gene diversity, expression, function, and evolution across vertebrates.

### Evolution of Contentious Phototransduction Recovery Gene Families in Frogs and Vertebrates

Although the roles and functions of the prominent genes and proteins involved in the vertebrate phototransduction cascade have been well-studied, there are still particular families and isoforms where debates and inconsistencies in both function and presence remain. In addition, research on the evolution of all genes and proteins involved in phototransduction both across and within vertebrate lineages is still lacking. In this study, we aimed to provide some clarity on the most contentious of these genes and proteins by determining if they are present in major vertebrate lineages and expressed in the eye, and by contextualizing these results in relation to the literature.

### Guanylyl Cyclases (GC-E, GC-F, and GC-D): Evidence for the Expression of an Olfactory GC in Frog Eyes

Guanylyl cyclases (GCs) are membrane-bound protein receptors found in several vertebrate systems that are responsible for signaling via cGMP (Kuhn, 2016). Among GCs, two are most commonly linked to the vertebrate visual system: GC-E encoded by *GUCY2D* (expressed in rods and cones) and GC-F encoded by *GUCY2F* (expressed in rods). These so-called visual GCs are responsible for synthesizing cGMP following activation by the guanylyl cyclase activating proteins (GCAPs) to restore the dark state current (*i.e.*, light-sensitive state) following photoreceptor excitation (Kuhn, 2016; Lamb & Hunt, 2018). Studies have shown that mutations in GC-E result in severe retinopathies (Perrault et al. 1996, 2000). In mice, mutations in GC-F cause severe cone dystrophy, but have little effect on rod cells (Yang et al. 1999). Knockout studies in mice targeting both of the visual GCs cause a complete lack of light-induced electrical responses and photoreceptor cell degeneration over time, highlighting the importance of these genes in the vertebrate visual system (Baehr et al. 2007). Across vertebrates there is variation in the number of GCs. *GUCY2D* is maintained in most vertebrate lineages while *GUCY2F* has been lost in reptiles and some mammals (*e.g.*, weasels, sloths, and marsupials) (Gesemann and Neuhauss, 2023).

Absence in some vertebrate lineages is thought to be linked to a nocturnal ancestor, creating a “bottleneck” effect where the gene is lost, and thus all descendant lineages also lack the gene (Gesemann and Neuhauss, 2023). However, we found that *GUCY2F* is found in lineages inferred to have had a nocturnal ancestry such as amphibians (including frogs) and mammals (lost in monotremes, and marsupials, but not placentals) indicating little support for this hypothesis as a universal explanation at a broad evolutionary scale. Alternatively, the decrease in number of GCs in some lineages may be related to a lack of need for a second visual guanylyl cyclase (Gesemann and Neuhauss, 2023).

Our results highlight the potential importance of another GC (*GC-D*) in the vertebrate visual system*. GC-D* is widespread across vertebrates and is typically referred to as the “olfactory guanylyl cyclase” due to its known function in the olfactory system (Kuhn, 2016) whereas there seems to exist confusion and lack of consensus surrounding this GC in the vertebrate visual system (Lamb & Hunt, 2018, Stiebel-Kalish et al. 2012). Previous studies have found evidence for *GC-D* in the genomes of Chondrichthyes, Coelacanthimorpha, Holostei, Teleostei, frogs, reptiles, and mammals (Lamb & Hunt, 2018; Lamb, 2020; Gesemann & Neuhauss, 2023), but fewer studies have provided evidence for expression of *GC-D* in the eye. Lamb and Hunt (2018) provided evidence for the expression of *GC-D* in the retinas of several Chondrichthyes and Holostei and reported that *GC-D* was the most highly expressed GC in the bowfin retina. They hypothesized that this GC may be important in aquatic environments, but further studies are needed to determine both the function of the “olfactory” GC and its potential relationship to aquatic environments. We found *GC-D* was present in all major jawed vertebrate lineages (including lungfish, salamanders, and caecilians, for which this had not previously been tested), but this is unsurprising given the role of this gene in the olfactory system. More importantly, we found evidence for eye expression of *GC-D* in the two species of teleost fishes that we examined, as well as in Chondrichthyes, amphibians (frogs), and reptiles (birds and snakes), suggesting a broader function of olfactory GC in vision across vertebrates. Further studies are needed, however, because many recovered transcripts were partial, and the number of species sampled was low, with strong evidence (complete transcripts) documented only in frogs, holost fishes, and one species of shark. The functional role of this GC in vision in frogs, fishes, and more broadly across vertebrates, as well as its cellular localization within the retina remain to be investigated.

### Guanylyl Cyclase Activating Proteins (GCAPs): Evidence for the Loss of GUCA1B2 Expression in Frog Eyes

Guanylyl Cyclase Activating Proteins (GCAPs) belong to a family of neuronal calcium sensor proteins and are responsible for regulating the activity of guanylyl cyclases (GCs) (*e.g.*, visual GC-E and GC-F, and olfactory GC-D discussed previously). There are three subfamilies of visual GCAPs; GCAP1 (encoded by *GUCA1A* and expressed mostly in cones), GCAP2 (encoded by *GUCA1B* and expressed mostly in rods) and GCAP3 (encoded by *GUCA1C* and expressed in cones) (Lamb, 2020). In addition, GCAP1 and GCAP2 each have two copies: GCAP1 (*GUCA1A*) and GCAPb (1-like) (*GUCA1A2*), and GCAP2 (*GUCA1B*) and GCAP2b (2-like) (*GUCA1B2*) (Gesemann and Neuhauss, 2023). Thus, in total there are five GCAPs that can be retained and expressed in vertebrates. Previous work on the function of GCAPs has focused on GCAP1, GCAP2, and GCAP3 with less work having been done on GCAP1L and GCAP2L (Lamb, 2020). The regulatory properties of GCAP1 and GCAP2 on the two visual GCs (GC-E and GC-F) are well known, and the remaining GCAPs serve similar functions with each GCAP having unique regulatory properties (Lamb, 2020). Consequently, there is variation in both the number of GCs and associated GCAPs amongst vertebrates. For example, highly visual species like many birds and lizards have retained only one GC but multiple GCAPs and thus maintain highly tunable regulation despite having only one GC (Gesemann and Neuhauss, 2023). On the other hand, some species adapted to dim-light conditions, such as blindsnakes and some nocturnal geckos, have fewer CGAPs, and thus reduced possibilities for tuning regulation (Gesemann and Neuhauss, 2023).

Gesemann and Neuhauss (2023) included five frog genomes in their study and found that four of the species retained all five GCAPs, but the direct-developing common coqui (*Eleutherodactylus coqui*) lacked a functional copy of *GUCA1B2* (GCAP2-b). Previous research has demonstrated that visual genes are expressed differentially across frog life-cycle stages (Schott et al. 2022a) and thus, Gesemann and Neuhauss (2023) proposed that the absence of *GUCA1B2* in the common coqui may be related to the life history of this species given that the other four frogs in their study are biphasic, with an aquatic larval stage. In our larger sample of frog species, *GUCA1B2* was the least frequently recovered gene, with a moderate recovery rate in the genomes (67%) and a very low recovery rate in the transcriptomes (18%). Additionally, more than half of the sequences of *GUCA1B2* we recovered from both the genomes and transcriptomes were partial and none of the transcriptomes from direct developing species showed evidence for the expression of this gene. The low recovery of *GUCA1B2*, however, extends beyond the direct-developing species in our dataset and so it seems that the loss, or loss of expression, is not explained by developmental mode. This was further corroborated by our transcriptome data from five species that included both tadpole and adult individuals. Among these, four demonstrated presence of *GUCA1B2* in both life stages, and only one (*Sclerophrys regularis*) had evidence for expression of *GUCA1B2* in the adult individual and not in the tadpole. Collectively, our results suggest that this gene, and protein, do not play an important, or even functional, role in the frog phototransduction pathway, and in species where it is retained it may serve a function outside the visual system. This contrasts with the pattern seen broadly across other vertebrates that retain *GUCA1B2* (several squamates and fishes) in which we found that transcriptomic evidence supported expression in the eye.

Finally, we also noted an interesting recovery pattern for *GUCA1C* (GCAP3) in frogs. While all GCAPs can regulate GC activity, there is variation in function. The visual GC, GC-E, has been shown to be predominantly regulated by GCAP1 in rods and Gesemann and Neuhauss (2023) suggested that as a result, GCAP2 and GCAP3 may be under lower evolutionary pressure and exhibit reduced expression. We recovered *GUCA1C* (GCAP3) in 82% of the genomes and 71% of the transcriptomes, with most sequences complete. Notably the recovery rate for *GUCA1C* was lower in both the genomes and transcriptomes than the other GCAPs: *GUCA1A*, *GUCA2A2*, and *GUCA1B*. We did not, however, find substantial differences in overall selective constraint acting on these genes, with overall selective constraint in *GUCA1C* (0.09215) being slightly higher than the other GCAPs (0.02165–0.05279), but close to the overall average for phototransduction genes (0.082509). Thus, despite expression of *GUCA1C* in eyes across frog species, we did not find substantial evidence to suggest *GUCA1C* is under lower evolutionary pressure compared to other GCAPs expressed in frog eyes.

### Guanylyl Cyclase Inhibiting Protein (GCIP): Maintenance and Expression in Vertebrates

In addition to the GCAPs, some vertebrates also retain a guanylyl cyclase inhibiting protein (GCIP) that is thought to serve a similar regulatory function in phototransduction, but the prevalence and specific function of this gene is not well understood (Lamb & Hunt, 2018; Lamb, 2020: Gesemann and Neuhauss, 2023). GCIP was first discovered by Li et al. (1998) while investigating GCAPs. In terms of function, GCIP is not able to stimulate GCs (unlike GCAPs) but does inhibit GC activity at high free calcium ion levels and competes with the GCAPs for the regulatory sites and thus function (Palczeski et al. 2004). In frog retinas, GCIP is localized to the inner segments, stomata, and the synaptic terminals of cone photoreceptors (Palczewski et al. 2004). In terms of evolutionary history, previous research has identified this protein only in frogs and teleosts and it has been noted as absent in many mammalian genomes (Palczewki et al. 2004). Evidence from Lamb (2020) supports the origin of GCIP as occurring before the whole-genome duplication in fishes and as the result of a duplication of the ancestral gene to GCIP/GCAP that occurred in the common ancestor of vertebrates, with inferred loss of GCIP in the last common ancestor of amniotes. We checked for the presence of GCIP in all other major vertebrate lineages and found it only in the genomes of non-amniotes, which broadly supports the hypotheses put forward by Lamb (2020), though broader surveys of GCIP expression in the eye are needed to investigate its potential role in phototransduction.

### Inhibitory Subunit of Phosphodiesterase (PDE6I): Limited Evidence in Frogs and Vertebrates More Broadly

Phosphodiesterase-6 (PDE6) plays a critical role in the light-activation phase of the vertebrate phototransduction cascade, but the activation of PDE6 is inhibited by the γ-subunit. As with many other vertebrate phototransduction genes, rod and cone photoreceptors possess cell-specific isoforms of both PDE6 and the inhibitory subunit (Wang et al. 2019). Specifically, among the inhibitory subunits, *PDE6G* is found in rods and *PDE6H* in cones (Lamb, 2020). A third inhibitory subunit of PDE6, *PDE6I*, was first described by Lagman et al. (2016), and additional evidence for the existence of the gene across vertebrates was provided by Lamb (2020). These studies found *PDE6I* sequences in Teleostei, Holostei, Coelacanthiformes, and Anura, but note that *PDE6I* was subsequently lost in amniotes. We found genomic evidence for putative *PDE6I* in additional lineages, namely Cladistia (bichir), Chondrostei (sturgeon and paddlefish), and Dipnoi (lungfish), as well as those previously identified but no others. Transcriptomic evidence for eye expression of *PDE6I* was previously limited to Florida gar (*Lepisosteus platyrhincus*) (Lamb et al. 2016), but we found eye expression of at least partial transcripts in one additional holost, a teleost, and 11 frog species. This low rate of recovery from the eye implies *PDE6I* does not play a role in the frog phototransduction cascade and possibly not in that of ray-finned fishes either. Our comparative analysis of the putative *PDE6I* sequences revealed a conserved 3’ region, but a highly variable 5’ region that had little to no homology across taxa. This suggests the possibility of erroneous gene predictions or that the sequences represent more than one gene. This later possibility was also supported by our phylogenetic analysis, which found multiple, and paraphyletic, *PDE6I* clades.

However, the short length of the conserved region of the gene makes phylogenetic inference unreliable (Lamb 2020). Further studies of comparative gene synteny and the tissue localization are needed to uncover the evolution and function of this enigmatic PDE gamma subunit.

### Surprisingly High Phototransduction Gene Diversity in Frogs

The last common ancestor of all jawed vertebrates was inferred to have had a complement of 38 phototransduction genes (not including the visual opsins or the putative *PDE6I*), with changes in this diversity over the course of vertebrate evolution (Lamb, 2020). The loss of phototransduction genes is often inferred to have been associated with a nocturnal evolutionary history, which is hypothesized to result from a reduced need for broad spectral sensitivity (Lamb & Hunt, 2018).

Mammals, for example, have experienced significant phototransduction gene loss linked to a nocturnal bottleneck early in their evolutionary history (Wu et al. 2017). Our results, however, suggest that frogs retain one of the most-complete repertoires of phototransduction genes among vertebrates. Only some ray-finned fish lineages retain more phototransduction genes. Nocturnality is inferred to be the ancestral state for frogs, and the majority of extant species are also primarily nocturnal (Anderson & Wiens, 2017); consequently, the retention of most phototransduction genes in frogs is in stark contrast to the pattern observed in other ancestrally nocturnal vertebrate lineages. They are consistent, however, with the retention of a more-complete repertoire of non-visual opsins in frogs relative to other ancestrally nocturnal lineages (Boyette et al. 2024).

The patterns of visual gene loss in squamates, birds, and mammals have alternatively been proposed to be linked to differences in photic habitat across life-cycle stages rather than nocturnal ancestry (Beaudry et al. 2017). Development in these groups takes place in dark environments (*i.e.*, shelled eggs or wombs), and thus perhaps it is these transitions from dark to light environments that are driving visual-gene evolution and diversity rather than nocturnal ancestry (Beaudry et al. 2017). Additionally, there is evidence for little visual gene loss in tuatara, an ancestrally nocturnal lineage, where maintenance of this diversity has been attributed to the usage of both bright-and dim-light environments across ontogeny (Gemmell et al. 2020). Similarly, many biphasic frogs are diurnal as tadpoles and nocturnal as adults (Ding et al. 2014; Altig & McDiarmid, 2015), but this difference is confounded with, in most species, a transition from an aquatic to terrestrial light environment that also imposes very different requirements on the visual system (Carlton et al. 2020; Schott et al. 2022a). Both factors have likely influenced the evolution of frog visual systems and contributed to their unique patterns.

### Positive Selection on Frog Phototransduction Genes Compared to other Vertebrates

Overall, we found evidence for positive selection acting on a subset of sites in a high proportion of frog phototransduction genes. Using the M8 PAML model we found 11 of the 36 phototransduction genes consistently expressed in frogs had significant evidence for positive selection. When accounting for synonymous rate variation using BUSTED, we identified over twice as many genes (25), nine of which overlap with those identified by M8. The two genes identified by M8 but not BUSTED may be false positives resulting from the lack of incorporation of synonymous rate variation with the M8 model. Sixteen genes were detected only when synonymous rate variation was accounted for using BUSTED, which suggests that BUSTED may be more sensitive than M8, perhaps due in part to the flexibility in site classes and their proportions, whereas M8 is restricted to the shape of a beta distribution. Given the differences in results between PAML and BUSTED, this highlights the importance of assessing multiple-model support in selection analyses.

We initially predicted greater evidence of positive selection in cone-specific phototransduction genes in frogs because of repeated transitions from nocturnality to diurnality. We found, however, that signals of positive selection were relatively evenly distributed across genes specific to cones and those found in both photoreceptor types (four genes each with evidence from both M8 and BUSTED). There was only limited evidence for positive selection in rod-specific phototransduction genes (one gene with evidence from both models), and genes involved in activation vs recovery also showed similar patterns (four genes each). In snakes, which also have several independent transitions to diurnality, 17 genes showed evidence of positive selection with most being cone-specific (Schott et al. 2018), a much stronger pattern than found in frogs. Birds have several activity pattern transitions, primarily in the opposite direction from diurnality to nocturnality, and evidence for positive selection has also been noted in a high proportion in the phototransduction genes (at least 14) (Wu et al. 2016; White et al. 2022). However, the positively selected genes in birds represent both cone-and rod-specific isoforms, as well as genes involved in activation and recovery, with no specific patterns identified. Among mammals, previous studies have shown evidence for positive selection for fewer genes (7), but interestingly, positive selection was restricted to rod-specific genes and those found in both photoreceptors (Wu et al. 2017). This suggests that patterns of positive selection among mammals are opposite to those found in snakes and distinct from the more general signals found in frogs and birds. Overall, these differences suggest considerable variation across vertebrates that is likely tied to the specific evolutionary and ecological histories of the various lineages.

In frogs, we found evidence for positive selection in both rod and cone receptor kinases (*GRK1* and *GRK7*). Receptor kinases play a critical role in the recovery phase of the vertebrate phototransduction cascade by phosphorylating the activated opsin protein (Fain et al. 2010).

Differential expression of both *GRK1* and *GRK7* between larval (tadpole) and juvenile southern leopard frogs, and between light and dark exposure conditions, is consistent with adaptive plasticity in receptor kinases in response to short-term changes in light levels and suggests these kinases could exhibit fixed functional adaptations to differing light environments (Schott et al. 2022). Combined with evidence for positive selection, this suggests these genes are undergoing functional tuning in frogs. Research has suggested that changes in the receptor kinases could speed up the phosphorylation of light-activated opsins, allowing for faster recovery in species that rapidly transition between bright and dark-light environments (Fu and Yau, 2007; Gurevich and Gurevich, 2010). In lineages of freshwater croakers (teleost fishes), alterations in RH1 from its ancestral state have been functionally linked to increased retinal release/recovery in rods, facilitating faster recovery for dim-light vision (Van Nynatten et al. 2021). Additionally, changes in the recovery protein arrestin in owls and deep-diving whales that specialize in dim-light environments with rapid bright-light transitions (and have rod-dominated retinas) are thought to help facilitate rapid sequestration of the phototoxic all-trans-retinal in rods, which would reduce the risk of light-induced retinopathy (Castiglione et al. 2023). Thus, alterations in the amino acid sequence of frog GRKs that result in faster recovery could be important in species that transition between aquatic and terrestrial environments, where the light environments vary and can change rapidly.

We also found evidence of positive selection in *SLC24A1*, which encodes one of the Na^+^/K^+^-Ca^2+^ ion exchangers found in vertebrates (Gurevich and Gurevich, 2010). These exchangers in combination with cyclic nucleotide gated channels (CNGs) are critical in regulating the current in photoreceptor cells, important in both the activation and recovery of the cells (Lamb, 2020).

Effective regulation of the movement of Ca^2+^ ions is important for triggering rapid recovery of the photoreceptors (Lamb, 2020). Essentially, Na^+^/K^+^-Ca^2+^ ion exchangers play a role in modulating the rate at which photoreceptors can recover and generate new responses (White et al. 2022). White et al. (2022) found evidence of positive selection in *SLC24A1* in nocturnal species (such as owls), which presumably have strong night vision, and manakins (diurnal passerine birds), which have been shown to have higher flicker fusion thresholds (high speed vision), and proposed that these patterns could be explained by adaptations for improved signal amplification and faster rod recovery, respectively. Similarly, Wu et al. (2016) found evidence of positive selection in *SLC24A1* in owls, which are primarily nocturnal, proposing a similar explanation for the results. Our results further indicate positive selection in two genes encoding CNGs (rod-specific *CNGA1* and cone-specific *CNGB3*) that also play a role in mediating recovery rates in photoreceptors (Lamb, 2020). Positive selection in *CNG*s has also been reported in some birds (owls and diurnal birds of prey), with the prediction that these changes could be involved in increased rod recovery rates in dim-light environments and may enhance the photoresponse (Wu et al. 2016). However, direct evidence linking changes in Na^+^/K^+^-Ca^2+^ ion exchangers or CNGs with enhanced visual capabilities has not been established.

### Signatures of Selection in Frog Phototransduction Genes Associated with Differences in Ecology

*Shifts in Adult Activity Pattern are not a Primary Driver of Frog Phototransduction Gene Evolution* We hypothesized that repeated transitions to diurnality from a nocturnal ancestry have driven adaptive visual evolution in diurnal frogs due to selective pressure for improved bright-light visual performance (*e.g.*, increases to temporal and spatial resolution). Based on this, we predicted evidence of positive selection in diurnal frog species, concentrated in cone-specific phototransduction genes, similar to the pattern found in snakes (Schott et al. 2018). Surprisingly, we did not find substantial evidence to support this. Of the eleven positively selected phototransduction genes identified by PAML, only two showed evidence for differences in selective constraint among diurnal species compared to non-diurnal species, the cone-specific phosphodiesterase *PDE6C* and “olfactory” guanylate cyclase *GC-D,* but the diurnal partition was not the best fit for these genes. That is, another trait better explained the differences in selective constraint in these genes rather than diurnality. BUSTED provided evidence for positive selection in twelve additional genes between diurnal and non-diurnal species, but this was not dominated by cone-specific genes. The signals of positive selection identified by BUSTED do not demonstrate a clear pattern between photoreceptor type or phototransduction phase. This fits into an emerging pattern that adult activity pattern is not a major driving force of visual evolution across frogs (Boyette et al. 2024, Schott et al. 2024), although lineage-specific forces may vary (Wan et al. 2023). There are, however, differences in activity pattern between life-cycle stages in many frog species, where nocturnal adult activity patterns are coupled with diurnal activity during the tadpole stage (Ding et al. 2014). This diurnal larval activity may result in visual systems being “preadapted” to diurnal conditions, lowering the strength of the selective pressure that shifts in activity patterns cause in other vertebrates. Broader sampling of frog species with different combinations of larval and adult activity patterns will be needed to test this hypothesis further.

### Scansorial Habitats are not Associated with Positive Selection in the Phototransduction Genes of Frogs

Based on morphological, spectral, and molecular evidence consistent with adaptation in scansorial frogs for increased spatial resolution (visual acuity) and temporal resolution to navigate complex, three-dimensional habitats, we hypothesized that scansoriality would be a strong predictor of positive selection for phototransduction genes. Overall, our results do not support this prediction, but we did find that significant differences in selective constraint between scansorial compared to non-scansorial frogs was the best explanation for differences in two genes, *GNAT2* and *RGS9BP3*.

*GNAT2* is the cone isoform of the alpha subunit of transducin and is responsible for activating the phosphodiesterase (Lamb et al. 2016). Interestingly, activated opsin proteins have been shown to activate alpha transducin at a constant rate, but the rate at which phosphodiesterase is activated is slower than transducin (Lamb et al. 2022). Changes in the alpha transducin could increase the rate at which phosphodiesterase is activated, and visual signals are subsequently communicated, which could be advantageous in complex arboreal environments, but this remains highly speculative. Possible explanations for the results in *RGS9BP3* are less clear. Only frogs and fishes maintain and express three copies of the RGS9 binding protein in the eye (Lamb et al. 2018). The third copy (*RGS9BP3*) is the least well studied among the three and its specific function in the eye is not known. Due to the limited information available, it is not clear how changes in this gene may be specific to scansorial species.

An alternative explanation for the pattern found in scansorial species could be related to the role gene expression levels play in alterations in protein function (Carlton et al. 2020).

Differences in expression levels of different phototransduction gene isoforms play a large role in activation and recovery speeds, and signal amplification, between the two photoreceptor cell types (Lamb & Hunt, 2018; Lamb, 2020). Evidence for differential expression has been documented in visual genes (including phototransduction genes) between tadpoles and juveniles of at least one frog species (Schott et al. 2022a). Thus, for *GNAT2* and *RGS9BP3,* differences in selection pressure in the absence of positive selection or significant relaxation or intensification could be explained by differences in gene expression levels between scansorial and non-scansorial species rather than direct changes in protein function.

### Geographic Distribution and Fossoriality are not Associated with Relaxed Selection in Frog Phototransduction Genes

Based on the known differences in photoperiod between tropical and temperate environments, we expected to see tropical frogs experience a relaxation of selective constraint, in particular in cone-specific phototransduction genes. Additionally, based on a divestment in the visual system by fossorial species, we also predicted evidence for a relaxation of selection, especially in cone genes. Overall, our results do not support these initial predictions. While we found that geographic distribution was the best explanation for differences in selective constraint for three genes (*CNGB3, GCIP, and GNB5*) what appears to be driving differences in selective constraint in these genes is either positive selection (*CNGB3*), the intensification of selection (*GNB5*), or neither (*GCIP*). Similarly, fossoriality was the best explanation for differences in selective constraint for only one phototransduction gene (*RGS9BP2*), but with no evidence for relaxed, intensified, or positive selection. The lack of a consistent pattern suggests that distribution, at least at this crude resolution, has not had a strong influence on phototransduction evolution in frogs. Previous research in both the visual opsins (Schott et al. 2024) and in non-visual opsins (Boyette et al. 2024) have addressed the importance of geographic distribution and fossoriality, but found that, despite the importance photoperiod and fossoriality for many aspects of frog physiology, their roles as a primary drivers of selective shifts in frog visual genes is not supported. However, we caution that small sample size for fossorial species is currently a limiting factor, and increased sampling will likely reveal the importance of this trait.

### Aquatic Environments and Differences in Life-History Strategy as Primary Drivers of Molecular Evolution in Frog Phototransduction Genes

Based on the spectral differences between aquatic and terrestrial environments and the role those differences have played on the evolution of visual genes both within frogs and across vertebrates, we expected to see differences in selective pressure between aquatic and terrestrial frog species and between direct-developing and non-direct developing species. We found that the direct-developing, fully aquatic, and semi-aquatic partitions best explain the patterns of selection seen in frog phototransduction genes, supporting our initial prediction. Combined, these three trait partitions were the best fit for 27 of the 37 phototransduction genes maintained in frogs (direct developing: ten genes, fully aquatic: eight genes, and fully + semi-aquatic: nine). We also found evidence for positive selection and relaxed selection in these genes across these partitions, which is discussed further below.

Direct-developing frogs are those without an aquatic tadpole stage where embryonic development is completed within the eggs, which hatch into miniature froglets (Kerney et al. 2010). Because direct-developing frogs lack aquatic tadpole stages, we expect selective pressures imposed on the visual system by aquatic light environments to be lacking in these frog species.

Schott et al (2022a) demonstrated differential expression of phototransduction genes across life-cycle stages in the southern leopard frog, a species with an aquatic tadpole stage, suggesting that visual genes are used differently during larval and adult stages. Here we found support for a release of selective constraint in direct developers based on significant evidence for relaxed selection acting on seven genes (*CNGB1*, *GRK7*, *GUCY2F*, *GC-D*, *PDE6A*, *PDE6B*, *SAG*). This relationship between life-history strategies and visual genes has also been found in non-visual opsins, where a relaxation of selection was identified in six genes between direct-developing and biphasic species (Boyette et al. 2024). With visual opsins, only one gene, the short wavelength sensitive 2 (*SWS2*) opsin, was found to show a difference for direct-developing species, where they were found to have significantly elevated *ω* that was inferred to be due to relaxed selection (Schott et al. 2024).

This gene was also found to be differentially expressed across life-cycle stages in the southern leopard frog (Schott et al. 2022a), which combined with evidence for relaxed constraint, suggests a difference in function or even importance in direct-developing species (Schott et al. 2024). Of the seven phototransduction genes with evidence for relaxed selection identified here, five were shown to be differentially expressed between tadpoles and juveniles, further supporting a potential functional shift in direct-developing species (Schott et al. 2022a).

Overall, the patterns seen across visual and other light-detection genes are consistent with the hypothesis that the visual and non-visual systems of direct-developing species have undergone a release in selective constraint due to functioning only in terrestrial light environments, although the effects are not equal across genes. *GUCA1A2* is an example of a gene that shows the opposite pattern. This gene was found to be under significant intensification of selection, but not positive selection (Table 1), which would suggest that *GUCA1A2* is under higher levels of selective constraint in direct-developing frogs than in biphasic species. A complete understanding of this vertebrate vision gene is still lacking. Previous studies have explored function and differences in expression of *GUCA1A* and *GUCA1B* in rod and cone photoreceptors (Cuenca et al. 1998).

However, the same level of attention has not been given to the other GCAPs, which had been attributed to the variable species-specific recovery patterns of the other genes (Gesemann & Neuhauss, 2023). Thus, although there is evidence for higher selective constraint in *GUCA1A2* in direct-developing frogs, a functional explanation remains unclear.

Beyond the patterns seen in direct-developing frogs, we see a strong pattern of the influence of aquatic environments on frog phototransduction genes. Previous studies have highlighted the impact of water on the transmission of light and there are numerous examples of visual-system adaptation to aquatic environments across vertebrates (*e.g.*, Schott et al. 2014; Van Nynatten et al. 2015; Hauser et al. 2021). We predict that frogs whose life history is intertwined with aquatic environments as adults are under greater selective pressure than those that are not associated with aquatic environments. The RELAX results potentially support this for some genes, where the fully aquatic and fully + semi-aquatic partitions also showed evidence for the intensification of selection (*RCVRN1*), positive selection (*GNB3, PDE6C, SLC24A2*), or both (*PDE6H, SLC24A1*). These results could suggest functional adaptations in fully aquatic species where these genes are under an intensification of selection. The long wavelength sensitive (*LWS*) opsin also exhibits evidence of relaxation of selective constraint in aquatic and fossorial species, suggesting these species may have undergone adaptations to the generally lower levels of available light found in these environments (Schott et al. 2024). This idea is also partially supported by the results of the direct-developing partitions, where it seems that lacking an aquatic life cycle stage does have a significant impact on the selective pressure exerted on the frog visual system.

*SLC24A1*, the gene that encodes Na^+^/K^+^-Ca^2+^ ion exchanger found in vertebrate rod photoreceptors, provides a potential example for evidence of functional adaptation of phototransduction driven by the aquatic environment. This ion exchanger, along with the cone-specific isoform encoded by *SLC24A2*, is important in modulating photoreceptor recovery rates (Gurevich and Gurevich, 2010; Lamb et al. 2020; White et al. 2022). *SLC24A1* demonstrates a signal of intensified selection in aquatic species that may reflect functional changes in this protein for a faster recovery time. Faster rod recovery time would increase response rates to new photons, which would be beneficial in lower-light aquatic environments. The pattern is less clear for the full + semi-aquatic partition where both species in our dataset that are fully aquatic were tested in combination with species that are semi-aquatic. We expected to see a similar pattern in the combined partition as in the fully aquatic partition, where there was evidence of potential functional adaptations to aquatic environments, but we found evidence for both relaxation of selection and positive selection in these genes. Given the variation in type and availability of light between aquatic and terrestrial environments, we would have predicted increased selective constraint (intensification) in these genes, because functional changes could move these genes, and their encoded proteins, away from a narrow functional optimum. Our results suggest that selection has actually relaxed on these genes, allowing them to change more easily. Given these findings, the functional implications of relaxed selection in these genes warrant further study to help clarify the role that light availability has played in visual-system evolution of aquatic frog species.

## Conclusions

The phototransduction cascade exemplifies critical sensory system processes in vertebrates that are also under selective pressure from the environment. Despite its importance, however, our understanding of the genes and proteins involved in phototransduction is still incomplete. We demonstrate that frogs retain among the most-complete repertoire of vertebrate phototransduction genes despite their nocturnal ancestry, and that (semi-)aquatic ecologies and major differences in developmental mode have been primary drivers of evolutionary change in frog phototransduction genes. Collectively, our results emphasize the important evolutionary gap that amphibians represent in our broader understanding of vision gene evolution as our vertebrate ancestors transitioned from living (and seeing) in water to living on land.

## Methods

### Phototransduction Gene Retention and Loss Across Vertebrates

There are numerous phototransduction genes that have been identified in vertebrates, but their retention and expression patterns across the major vertebrate lineages are not fully clear. In order to contextualize broad patterns of phototransduction gene evolution across frogs we first identified known patterns of gene retention, loss, and duplication across major vertebrate lineages using literature searches. We then identified possible query sequences for all vertebrate phototransduction genes and used targeted BLAST (Altschul et al. 1990) searches against the NCBI core nucleotide dataset to check for presence or absence in the jawed vertebrate lineages outlined in Figure 2. We then identified representative eye transcriptomes for further examination. A full list of the transcriptomes used for these analyses can be found in Supplementary File S1. We focused expression analyses on four genes that have not been well characterized: “olfactory” guanylyl cyclase (*GC-D*), guanylyl cyclase activating protein 1 B2 (*GUCA1B2*), guanylyl cyclase inhibiting protein (*GCIP*), and an isoform of the inhibitory subunit of phosphodiesterase (*PDE6I*). We used both published sequences and those from the present study as references for BLAST and Genbank searches. We recorded the presence and absence of genes in the genomes and expression in the transcriptomes following the criteria outlined below. Phylogenetic relationships among vertebrate lineages were described following Irisarri et al (2017).

Further analysis was conducted on *PDE6I* because prior studies (Lagman et al. 2016; Lamb 2020) have not conducted a phylogenetic analysis on this gene and its distribution and function across vertebrates is unclear. Starting from sequences identified in previous studies (Lagman et al. 2016; Lamb 2020), we searched for PDE6 gamma genes (*PDE6G*, *PDE6H*, and *PDE6I*) using the approach outlined above but also including agnathans. We extracted a representative sample of *PDE6G* and *PDE6H* from NCBI for these lineages, and all putative *PDE6I* sequences we identified in the nucleotide database and the eye transcriptomes from frogs and the other vertebrates. The three sets of sequences were aligned together using MACSE (Ranwez et al. 2018) and trimmed using ClipKIT smart-gap trimming and codon mode (Steenwyk et al. 2020). Inspection of the alignment revealed a lack of homology across taxa 5’ of the start codon of *PDE6G* and *PDE6H* genes. This region was manually trimmed followed by an additional round of MACSE alignment and two rounds of ClipKit trimming. A maximum likelihood (ML) gene tree was inferred using PhyML 3 (Guindon et al. 2010) under the GTR + G + I nucleotide model with five rate categories, empirical nucleotide frequencies, maximum likelihood estimated transition to transversion rate ratios, and SH-like branch support values. In the absence of a more appropriate outgroup, the tree was re-rooted on the branch leading to agnathan *PDE6G*.

### Frog Species Sampling

We utilized frog eye transcriptomes generated by previous studies (Schott et al. 2022, 2024). NCBI’s Genbank and Genome databases were also searched for annotated sequences and genome assemblies to increase the phylogenetic and ecological diversity of the dataset. Our combined sampling efforts include 113 species, representing 34 of 57 currently recognized (*e.g.*, AmphibiaWeb, 2025) frog families, including 4 species with both transcriptomic and genomic data and 6 species with both tadpole and adult transcriptomic data (Fig. 2; Schott et al. 2022, 2024).

### Phototransduction Sequence Recovery in Frogs

Phototransduction gene coding sequences were extracted from frog genomes using vertebrate query sequences obtained previously (Schott et al. 2018; Gemmell et al. 2020) using BLAST and Genbank searches (Altschul et al. 1990). These were subsequently used as query sequences for the remaining frog genomes and eye transcriptomes. Recovered phototransduction gene sequences were manually assembled and aligned against the query sequences in MEGA (Tamura et al. 2013). A coding sequence was considered complete if it was recovered entirely from start to stop codon. Partial sequences were classified as those sequences where 50%–99% of the codons were recovered. Sequences with less than 50% of the codons recovered were considered incomplete and were excluded from analyses but were differentiated from sequences that were completely absent (not recovered). Any premature stop codons were converted to gaps to enable inclusion in downstream analyses (Yohe et al. 2017). Genes were considered expressed in frog eyes if they were completely or partially recovered (>50%) in the transcriptomes. Prior to further analysis, the gene identity of each recovered sequence was confirmed by phylogenetic analysis. Any genes that were misidentified were reextracted and if the correct gene could still not be recovered, were considered absent. A full list of genes recovered from the frogs used in this study can be found in Supplementary File S2, online supplementary material.

### Frog Trait Classification

Frog species were partitioned into discrete categories for each of seven ecology or life-history traits that we predicted to influence the evolution of phototransduction genes including: diurnal/nocturnal, fully aquatic/terrestrial, fully + semi-aquatic/terrestrial, fossorial/non-fossorial, scansorial/non-scansorial, tropical/temperate, and direct-developing/non-direct-developing based on previous studies (Thomas et al. 2020; Boyette et al. 2024; Schott et al. 2024).

### Selection Analyses

Coding regions for each of the recovered phototransduction genes were aligned with MACSE (Ranwez et al. 2018) and PRANK (Lötytnoja, 2014) alignment tools through an in-house pipeline, followed by manual correction, as necessary, to correct mis-assemblies, or to remove nonhomologous sequence regions that can result from automated gene prediction (*e.g.*, intronic sequence). ML gene trees were inferred using PhyML 3 (Guindon et al. 2010) under the GTR + G + I nucleotide model with five rate categories, empirical nucleotide frequencies, and maximum likelihood estimated transition to transversion rate ratios. Because individual gene trees do not always reflect the evolutionary history of a species, it is best practice to compare results for selection analyses using both gene-and species-tree topologies (*e.g.*, Schott et al. 2018, 2024; Van Nynatten et al. 2021). This ensures results are robust to minor topological differences. All analyses in this study were performed using both the ML gene tree and a species-tree topology based on the phylogeny of Jetz and Pyron (2018). Species topologies were generated by editing PhyML gene topologies using an inhouse pipeline and the ‘drop.tip’ function in the R (v4.4.1) package ‘ape’ (v5.8; Paradis and Schliep, 2019). All topologies were modified to contain the basal trichotomy required by PAML (Yang, 2007) and pruned to match the taxa in each alignment for each of the 37 phototransduction genes tested.

Each tree was paired with its respective gene alignment and analyzed using the PAML 4 random site models (M0, M1a, M2a, M2a_rel, M3, M7, M8a, and M8) to estimate the rates of nonsynonymous to synonymous nucleotide substitutions (⍵ or d*N*/d*S*) (Yang, 2007) using the BlastPhyMe interface (Schott et al. 2016). This allowed for the inference of alignment-wide selection patterns and to test for positive selection. All analyses were run using varying starting values to avoid potential local optima. The significance and best-fit among model pairs was determined using a likelihood ratio test (LRT) with a *χ*^2^ distribution. Evidence for relationships between photoreceptor cell type or phototransduction phase was evaluated using a Mann-Whitney U test in R.

To test if shifts in selection among frog phototransduction genes corresponded to variation in life history or ecology, we used the PAML clade model C (Bielawski and Yang, 2004). This model enables tests for evidence of a codon site class that demonstrates a shift in selection between the pre-partitioned foreground and background groups (*e.g.*, diurnal vs non-diurnal frogs). Any combination of branches or clades within a phylogeny can form these groups. The results from CmC (assumes a class of sites is free to evolve differently among two or more partitions; *ω*_D1_ > 0 and *ω*_D1_ ≠ *ω*_D2_ > 0) are compared to the null model M2a_rel.

In cases where PAML analyses found evidence for significantly elevated *ω* in a group of interest (*e.g.*, elevated ω in diurnal frogs vs non-diurnal frogs), we used the models BUSTED (Murrell et al. 2015) and RELAX (Wertheim et al. 2015) implemented on the Datamonkey server (Delport et al. 2010). BUSTED provides an alternative approach to identifying episodic positive selection across a gene. If used on an entire phylogeny, the results are comparable to the PAML M2a model or, if used on separate partitions of a phylogeny, results are comparable to PAML clade models. We performed BUSTED analyses using both approaches. BUSTED differs from PAML in the inclusion of synonymous rate variation, providing no specified neutral site class, and does not constrict the proportion of sites in each site class to be the same between the null and alternative models. The RELAX model is designed to test whether an elevated ω is likely to be the result of relaxed selective constraint (*i.e.*, a lack of selection against change) or adaptive selection (*i.e.*, selection for change). A value of K<1 indicates relaxed selection on the foreground branches compared to the background and when K>1 this indicates adaptive selection on the foreground group compared to the background group. Given the number of genes with multiple models being tested, all positive results were also corrected for false discovery rate (q<0.05) under the Benjamini-Hochberg procedure using a custom script.

## Data Availability

The data underlying this article are available on NCBI under Bioproject PRJNA1073881 and on Zenodo (Perez et al. 2025; Schott et al. 2024b), as well as in the Supplementary Materials. Scripts used for data analysis are available on GitHub (https://github.com/Schott-Lab/).

## Authors’ Contributions

RKS conceived and designed the study, with input from RCB, MKF, KNT, DJG, and JWS. RCB, MKF, DJG, and JWS conducted fieldwork. TJMP, RKS, and CL extracted the gene sequences and performed the molecular evolutionary analyses. TJMP, RKS, and RCB wrote the manuscript. RKS provided supervision. RKS, RCB, DJG, JWS, and MKF provided project administration and acquired funding. All authors reviewed and editted the final manuscript.

## Acknowledgements

We thank Amirmohammad Nasiri, Aaiza Khan, Renée Gorman, Sabrina Brusco, and Phoebe Hall for their assistance curating the dataset used in this study. We thank the following field companions who helped obtain specimens for this work: Hannah Augustijnen, Abraham G. Bamba Kaya, C. Guillherme Becker, Gabriela Bittencourt-Silva, Itzue Calviedes Solis, Patrick Campbell, Diego Cisneros-Heredia, Simon Clulow, Christian L. Cox, Mateo Davila, Paul Doughty, Juvencio Eko Mengue, TJ Firneno, Carl Franklin, Philippe Gaucher, Ivan Gomez-Mestre, Shakuntala Devi Gopal, Jon and Krittee Gower, Célio F. B. Haddad, Anthony Herrel, Sunita Janssenswillen, Jim Labisko, H. Christoph Liedtke, Simon Loader, Simon Maddock, Michael Mahony, Renato A. Martins, Matthew McElroy, Christopher Michaels, Nicki Mitchell, Justino Nguema Mituy, Diego Moura, Martin Nsue, Daniel M. Portik, Ivan Prates, Kim Roelants, Corey Roelke, Lauren Scheinberg, Bruno Simões, Ben Tapley, Elie Tobi, Rose Upton, Mark Wilkinson, and Molly Womack. We thank the Gabon Biodiversity Program and Bioko Biodiversity Protection Program for logistical support in the field; Grant Webster, Scott Keogh, and Jared Grummer for advice on where to find key species; Carolina Reyes-Puig for help with specimen numbers; and Jodi Rowley and Stephen Mahony for assistance exporting tissues for analysis. Sampling was conducted following IACUC protocols (NHMUK, NMNH 2016-012, UNESP Rio Claro CEUA-23/ 2017, UTA A17.005, ANU A2017/47) and with scientific research authorizations (the United States: Texas Parks and Wildlife Division SR-0814-159, North Cascades National Parks NCCO-2018-SCI-0009; Brazil: ICMBio MMA 22511-4, ICMBio SISBIO 30309-12; the United Kingdom: NE Licence WML-OR04; French Guiana: RAA: R03-2018-06-12-006; Gabon: CENAREST AR0020/17; Australia: New South Wales National Parks & Wildlife Service SL102014, Queensland Department of National Parks WITK18705517; Equatorial Guinea: INDEFOR-AP 0130/020-2019). This research was supported by an NSERC Discovery Grant (to R.K.S.) and grants from the Natural Environment Research Council, UK (NE/R002150/1) and the National Science Foundation (DEB-1655751).

## Notes

### Competing Interest Statement

The authors have declared no competing interest.

## References

Altschul SF, Gish W, Miller W, Myers EW, Lipman DJ. 1990. Basic Local Alignment Search Tool. Journal of Molecular Biology. 215:403–410.

Altig R, McDiarmid R. 2015. Handbook of larval amphibians of the United States and Canada. Ithaca: Comstock Publishing Associates.

AmphibiaWeb. 2025. <https://amphibiaweb.org> University of California, Berkeley, CA, USA

Anderson SR, Wiens JJ. 2017. Out of the dark: 350 million years of conservatism and evolution in diel activity patterns in vertebrates: EVOLUTION OF DAY-NIGHT ACTIVITY PATTERNS. Evolution. 71(8):1944–1959. doi:10.1111/evo.13284.

Arinobu D, Tachibanaki S, Kawamura S. 2010. Larger inhibition of visual pigment kinase in cones than in rods. Journal of Neurochemistry. 115(1):259–268. doi:10.1111/j.1471-4159.2010.06925.x.

Arshavsky, V. Y., and Burns, M. E. (2012). Photoreceptor signaling: Supporting vision across a wide range of light intensities. Journal of Biological Chemistry, 287(3), 1620–1626. 10.1074/jbc.R111.305243.

Baehr W, Karan S, Maeda T, Luo D-G, Li S, Bronson JD, Watt CB, Yau K-W, Frederick JM, Palczewski K. 2007. The Function of Guanylate Cyclase 1 and Guanylate Cyclase 2 in Rod and Cone Photoreceptors. Journal of Biological Chemistry. 282(12):8837–8847. doi:10.1074/jbc.M610369200.

Beaudry FEG, Iwanicki TW, Mariluz BRZ, Darnet S, Brinkmann H, Schneider P, Taylor JS. 2017. The non-visual opsins: eighteen in the ancestor of vertebrates, astonishing increase in ray-finned fish, and loss in amniotes. J Exp Zool Pt B. 328(7):685–696. doi:10.1002/jez.b.22773.

Bielawski JP, Yang Z. 2004. A Maximum Likelihood Method for Detecting Functional Divergence at Individual Codon Sites, with Application to Gene Family Evolution. J Mol Evol. 59(1). doi:10.1007/s00239-004-2597-8.

Borah BK, Renthlei Z, Trivedi AK. 2019. Seasonality in terai tree frog (Polypedates teraiensis): Role of light and temperature in regulation of seasonal breeding. Journal of Photochemistry and Photobiology B: Biology. 191:44–51. 10.1016/j.jphotobiol.2018.12.005

Bowmaker JK. 2008. Evolution of vertebrate visual pigments. Vision Research. 48(20):2022–2041. doi:10.1016/j.visres.2008.03.025.

Boyette JL, Bell RC, Fujita MK, Thomas KN, Streicher JW, Gower DJ, Schott RK. (2024). Diversity and molecular evolution of non-visual opsin genes across environmental, developmental, and morphological adaptations in frogs. Molecular Biology and Evolution.:msae090. doi:10.1093/molbev/msae090.

Buchanan, B. (2006). Observed and potential effects of artificial night lighting on frog amphibians. In C. Rich, & T. Longcore (Eds.), Ecological consequences of artificial night lighting (pp. 192–220). Island Press.

Burns, M. E., and Pugh, E. N., Jr. (2010). Lessons from photoreceptors: Turning off G-protein signaling in living cells. Physiology, 25(2), 72–84. 10.1152/physiol.00001.2010

Carleton KL, Escobar-Camacho D, Stieb SM, Cortesi F, Marshall NJ. 2020. Seeing the rainbow: mechanisms underlying spectral sensitivity in teleost fishes. Journal of Experimental Biology. 223(8):jeb193334. doi:10.1242/jeb.193334.

Castiglione GM, Chiu YLI, Gutierrez E de A, Van Nynatten A, Hauser FE, Preston M, Bhattacharyya N, Schott RK, Chang BSW. 2023. Convergent evolution of dim light vision in owls and deep-diving whales. Current Biology. 33:1–8.10.1016/j.cub.2023.09.015

Castoe TA, De Koning APJ, Hall KT, Card DC, Schield DR, Fujita MK, Ruggiero RP, Degner JF, Daza JM, Gu W, et al. 2013. The Burmese python genome reveals the molecular basis for extreme adaptation in snakes. Proc Natl Acad Sci USA. 110(51):20645–20650. doi:10.1073/pnas.1314475110.

Corbo JC. 2021. Vitamin A1/A2 chromophore exchange: Its role in spectral tuning and visual plasticity. Developmental Biology. 475:145–155. doi:10.1016/j.ydbio.2021.03.002.

Cronin TW, Johnsen S, Marshall J, Warrant EJ. Visual ecology. Princeton: Princeton University Press; 2014.

Davies WIL, Collin SP, Hunt DM. 2012. Molecular ecology and adaptation of visual photopigments in craniates. Molecular Ecology. 21(13):3121–3158. doi:10.1111/j.1365-294X.2012.05617.x.

Davies WIL, Tamai TK, Zheng L, Fu JK, Rihel J, Foster RG, Whitmore D, Hankins MW. 2015. An extended family of novel vertebrate photopigments is widely expressed and displays a diversity of function. Genome Res. 25(11):1666–1679. doi:10.1101/gr.189886.115.

Delport W, Poon AFY, Frost SDW, Kosakovsky Pond SL. 2010. Datamonkey 2010: a suite of phylogenetic analysis tools for evolutionary biology. Bioinformatics. 26(19):2455–2457. doi:10.1093/bioinformatics/btq429.

Ding G-H et al. 2014. Effects of light intensity on activity in four sympatric anuran tadpoles. Zoological Research. 35(4). doi:10.13918/j.issn.2095-8137.2014.4.332

Fain GL, Hardie R, Laughlin SB. 2010. Phototransduction and the Evolution of Photoreceptors. Current Biology. 20(3):R114–R124. doi:10.1016/j.cub.2009.12.006.

Fu Y, Yau K-W. 2007. Phototransduction in mouse rods and cones. Pflugers Arch - Eur J Physiol. 454(5):805–819. doi:10.1007/s00424-006-0194-y.

Gemmelll NJ, Rutherford K, Prost S, Tollis M, Winter D, Macey JR, Adelson DL, Suh A, Bertozzi T, Grau JH, et al. 2020. The tuatara genome reveals ancient features of amniote evolution. Nature. 584(7821):403–409. doi:10.1038/s41586-020-2561-9.

Gesemann M, Neuhauss SCF. 2023. Evolution of visual guanylyl cyclases and their activating proteins with respect to clade and species-specific visual system adaptation. Front Mol Neurosci. 16:1131093. doi:10.3389/fnmol.2023.1131093.

Guindon S, Dufayard JF, Lefort V, Anisimova M, Hordijk W, Gascuel O. New algorithms and methods to estimate maximumlikelihood phylogenies: assessing the performance of PhyML 3.0. Syst Biol. 2010:59(3):307–321. 10.1093/sysbio/syq010.

Gurevich VV, Gurevich EV. 2010. Phototransduction: Inactivation in Cones. Elsevier Ltd. pg: 397–402.

Hauser FE, Ilves KL, Schott RK, Alvi E, López-Fernández H, Chang BSW. 2021. Evolution, inactivation and loss of short wavelength-sensitive opsin genes during the diversification of Neotropical cichlids. Mol Ecol. 30(7):1688–1703. doi:10.1111/mec.15838.

Hauser FE, Chang BS. 2017. Insights into visual pigment adaptation and diversity from model ecological and evolutionary systems. Current Opinion in Genetics & Development. 47:110–120. doi:10.1016/j.gde.2017.09.005.

Helten A, Säftel W, Koch K. 2007. Expression level and activity profile of membrane bound guanylate cyclase type 2 in rod outer segments. Journal of Neurochemistry. 103(4):1439–1446. doi:10.1111/j.1471-4159.2007.04923.x.

Huang CH, Zhong MJ, Liao WB, Kotrschal A. 2019. Investigating the role of body size, ecology, and behavior in frog eye size evolution. Evol Ecol. 33(4):585–598. doi:10.1007/s10682-019-09993-0.

Ingram NT, Sampath AP, Fain GL. 2016. Why are rods more sensitive than cones? The Journal of Physiology. 594(19):5415–5426. doi:10.1113/JP272556.

Irisarri I, Baurain D, Brinkmann H, Delsuc F., et al. 2017. Phylotranscriptomic consolidation of the jawed vertebrate timetree. Nat Ecol Evol. 1(9):1370–1378. 10.1038/s41559-017-0240-5

IUCN. 2022. The IUCN Red List of Threatened Species. Version 2022-1.https://www.iucnredlist.org. Accessed on [23 September 2022].

Jetz W, Pyron RA. 2018. The interplay of past diversification and evolutionary isolation with present imperilment across the amphibian tree of life. Nat Ecol Evol. 2(5):850–858. doi:10.1038/s41559-018-0515-5.

Kawamura S, Tachibanaki S. 2022. Molecular bases of rod and cone differences. Progress in Retinal and Eye Research. 90:101040. doi:10.1016/j.preteyeres.2021.101040.

Kerney R, Gross JB, Hanken J. 2010. Early cranial patterning in the direct-developing frog *Eleutherodactylus coqui* revealed through gene expression: Early cranial patterning in *E. coqui*. Evolution & Development. 12(4):373–382. doi:10.1111/j.1525-142X.2010.00424.x.

Kefalov, V. J. (2012). Rod and cone visual pigments and phototransduction through pharmacological, genetic, and physiological approaches. Journal of Biological Chemistry, 287(3), 1635–1641. 10.1074/jbc.R111.303008

Kojima K, Matsutani Y, Yamashita T, Yanagawa M, Imamoto Y, Yamano Y, Wada A, Hisatomi O, Nishikawa K, Sakurai K, et al. 2017. Adaptation of cone pigments found in green rods for scotopic vision through a single amino acid mutation. Proc Natl Acad Sci USA. 114(21):5437–5442. doi:10.1073/pnas.1620010114.

Koskelainen A, Hemilä S, Donner K. 1994. Spectral sensitivities of short-and long-wavelength sensitive cone mechanisms in the frog retina. Acta Physiologica Scandinavica. 152(1):115–124. doi:10.1111/j.1748-1716.1994.tb09790.x.

Kuhn M. 2016. Molecular Physiology of Membrane Guanylyl Cyclase Receptors. Physiological Reviews. 96(2):751–804. doi:10.1152/physrev.00022.2015.

Lamb TD. 2016. Why rods and cones? Eye. 30(2):179–185. doi:10.1038/eye.2015.236.

Lamb TD, Patel H, Chuah A, Natoli RC, Davies WIL, Hart NS, Collin SP, Hunt DM. 2016. Evolution of Vertebrate Phototransduction: Cascade Activation. Mol Biol Evol. 33(8):2064–2087. doi:10.1093/molbev/msw095.

Lamb TD, Hunt DM. 2018. Evolution of the calcium feedback steps of vertebrate phototransduction. Open Biol. 8(9):180119. doi:10.1098/rsob.180119.

Lamb TD. 2020. Evolution of the genes mediating phototransduction in rod and cone photoreceptors. Progress in Retinal and Eye Research. 76:100823. doi:10.1016/j.preteyeres.2019.100823.

Lamb TD. 2022. Photoreceptor physiology and evolution: cellular and molecular basis of rod and cone phototransduction. The Journal of Physiology. 600(21):4585–4601. doi:10.1113/JP282058.

Larhammar D, Nordström K, Larsson TA. 2009. Evolution of vertebrate rod and cone phototransduction genes. Philos Trans R Soc Lond B Biol Sci. 364(1531):2867–2880. doi:10.1098/rstb.2009.0077.

Li N, Fariss R, Zhang K, Otto-Bruc A, Haeseleer F, Bronson JD, Qin N, Yamazaki A, Subbaraya I, Milam AH, et al. 1998. Guanylate-cyclase-inhibitory protein is a frog retinal Ca2-binding protein related to mammalian guanylate cyclase activating proteins. European Journal of Biochemistry. 252:591–599.

Löytynoja A “Phylogeny-aware alignment with PRANK.” Methods Mol. Biol. 2014;1079 10.1007/978-1-62703-646-7_10.

Macpherson ESB, Hauser FE, Van Nynatten A, Chang BSW, Lovejoy NR. 2024. Evolution of rhodopsin in flatfishes (Pleuronectiformes) is associated with depth and migratory behavior. Journal of Fish Biology. 105(3):779–790. doi:10.1111/jfb.15828.

Makino CL, Dodd RL, Chen J, Burns ME, Roca A, Simon MI, Baylor DA. 2004. Recoverin Regulates Light-dependent Phosphodiesterase Activity in Retinal Rods. The Journal of General Physiology. 123(6):729–741. doi:10.1085/jgp.200308994.

Mitra AT, Womack MC, Gower DJ, Streicher JW, Clark B, Bell RC, Schott RK, Fujita MK, Thomas KN. 2022. Ocular lens morphology is influenced by ecology and metamorphosis in frogs and toads. Proc R Soc B. 289(1987):20220767. doi:10.1098/rspb.2022.0767.

Mohun SM, Davies WL, Bowmaker JK, Pisani D, Himstedt W, Gower DJ, Hunt DM, Wilkinson M. 2010. Identification and characterization of visual pigments in caecilians (Amphibia: Gymnophiona), an order of limbless vertebrates with rudimentary eyes. Journal of Experimental Biology. 213(20):3586–3592. doi:10.1242/jeb.045914.

Mohun SM, Wilkinson M. 2015. The eye of the caecilian *Rhinatrema bivittatum* (Amphibia: Gymnophiona: Rhinatrematidae). Acta Zoologica. 96(2):147–153. doi:10.1111/azo.12061.

Murrell B, Moola S, Mabona A, Weighill T, Sheward D, Kosakovsky Pond SL, Scheffler K. 2013. FUBAR: A Fast, Unconstrained Bayesian AppRoximation for Inferring Selection. Molecular Biology and Evolution. 30(5):1196–1205. doi:10.1093/molbev/mst030.

Murrell B, Weaver S, Smith MD, Wertheim JO, Murrell S, Aylward A, Eren K, Pollner T, Martin DP, Smith DM, et al. 2015. Gene-Wide Identification of Episodic Selection. Molecular Biology and Evolution. 32(5):1365–1371. doi:10.1093/molbev/msv035.

Musilova Z, Cortesi F, Matschiner M, Davies WIL, Patel JS, Stieb SM, de Busserolles F, Malmstrøm M, Tørresen OK, Brown CJ, et al. 2019. Vision using multiple distinct rod opsins in deep-sea fishes. Science. 364: 588–592.

Navarrete Méndez MJ, Amini SS, Santos JC, Saal J, Wake MH, Ron SR, Tarvin RD. 2025. Caecilians maintain a functional long-wavelength-sensitive cone opsin gene despite signatures of relaxed selection and more than 200 million years of fossoriality. doi:10.1101/2025.02.07.636964. [accessed 2025 Feb 18]. http://biorxiv.org/lookup/doi/10.1101/2025.02.07.636964.

Nilsson D-E. 2021. The Diversity of Eyes and Vision. Annu Rev Vis Sci. 7(1):19–41. doi:10.1146/annurev-vision-121820-074736.

Ozawa T, Fukuda M, Nara M, Nakamura A, Komine Y, Kohama K, Umezawa Y. 2000. How Can Ca^2+^ Selectively Activate Recoverin in the Presence of Mg^2+^Surface Plasmon Resonance and FT-IR Spectroscopic Studies. Biochemistry. 39(47):14495–14503. doi:10.1021/bi001930y.

Palczewski K. 2014. Chemistry and Biology of the Initial Steps in Vision: The Friedenwald Lecture. Invest Ophthalmol Vis Sci. 55(10):6651. doi:10.1167/iovs.14-15502.

Palczewski K, Sokal I, Baehr W. 2004. Guanylate cyclase-activating proteins: structure, function, and diversity. Biochemical and Biophysical Research Communications. 322(4):1123–1130. doi:10.1016/j.bbrc.2004.07.122.

Paradis E and Schliep K. 2019. “ape 5.0: an environment for modern phylogenetics and evolutionary analyses in R.” Bioinformatics, 35, 526–528. doi:10.1093/bioinformatics/bty633.

Perez, T., Bell, R., Lavoie, C., Fujita, M., Gower, D. J., Streicher, J., Thomas, K. N., & Schott, R. 2025. Data from: Evolutionary and ontogenetic shifts from aquatic to terrestrial light environments in frogs provide new insights into the vertebrate phototransduction cascade [Data set]. Zenodo. 10.5281/zenodo.17487275

Perrault I et al. 1996 Retinal-specific guanylate cyclase gene mutations in Leber’s congenital amaurosis. Nat. Genet. 14, 461 –464. (doi:10.1038/ ng1296-461)

Perrault I, Rozet JM, Gerber S, Ghazi I, Ducroq D, Souied E, Leowski C, Bonnemaison M, Dufier JL, Munnich A, Kaplan J. 2000. Spectrum of retGC1 mutations in Leber’s congenital amaurosis. European Journal of Human Genetics. Aug;8(8):578–82.

Ranwez V, Harispe S, Delsuc F, Douzery EJP. 2011. MACSE: Multiple Alignment of Coding SEquences Accounting for Frameshifts and Stop Codons. Murphy WJ, editor. PLoS ONE. 6(9):e22594. doi:10.1371/journal.pone.0022594.

Ranwez V, Douzery EJP, Cambon C, Chantret N, Delsuc F. 2018. MACSE v2: Toolkit for the Alignment of Coding Sequences Accounting for Frameshifts and Stop Codons. Molecular Biology and Evolution. 35(10):2582–2584. doi:10.1093/molbev/msy159.

Schott RK, Gow D, Chang BS. 2016a. BlastPhyMe: A toolkit for rapid generation and analysis of protein-coding sequence datasets. Bioinformatics. [accessed 2024 Mar 3]. http://biorxiv.org/lookup/doi/10.1101/059881.

Schott RK, Van Nynatten A, Card DC, Castoe TA, Chang BSW. 2018. Shifts in Selective Pressures on Snake Phototransduction Genes Associated with Photoreceptor Transmutation and Dim-Light Ancestry. Molecular Biology and Evolution. 35(6):1376–1389. doi:10.1093/molbev/msy025.

Schott RK, Bhattacharyya N, Chang BSW. 2019. Evolutionary signatures of photoreceptor transmutation in geckos reveal potential adaptation and convergence with snakes. Evolution. 73(9):1958–1971. doi:10.1111/evo.13810.

Schott RK, Bell RC, Loew ER, Thomas KN, Gower DJ, Streicher JW, Fujita MK. 2022a. Transcriptomic evidence for visual adaptation during the aquatic to terrestrial metamorphosis in leopard frogs. BMC Biol. 20(1):138. doi:10.1186/s12915-022-01341-z.

Schott RK, Perez L, Kwiatkowski MA, Imhoff V, Gumm JM. 2022b. Evolutionary analyses of visual opsin genes in frogs and toads: Diversity, duplication, and positive selection. Ecology and Evolution 12(2). doi:10.1002/ece3.8595.

Schott RK, Fujita MK, Streicher JW, Gower DJ, Thomas KN, Loew ER, Bamba Kaya AG, Bittencourt-Silva GB, Guillherme Becker C, Cisneros-Heredia D, et al. 2024. Diversity and Evolution of Frog Visual Opsins: Spectral Tuning and Adaptation to Distinct Light Environments. Molecular Biology and Evolution. 41(4):msae049. doi:10.1093/molbev/msae049.

Schott, R., Fujita, M., Streicher, J., Gower, D., Thomas, K., & Bell, R. 2024b. Frog Eye Transriptome Assemblies [Data set]. In Molecular Biology and Evolution (Vol. 41, Number 4, p. msae049). Zenodo. 10.5281/zenodo.12535262

Steenwyk JL, Buida TJ III, Li Y, Shen X-X, Rokas A. 2020. ClipKIT: A multiple sequence alignment trimming software for accurate phylogenomic inference. PLOS Biology 18:e3001007.

Shrimpton SJ, Streicher JW, Gower DJ, Bell RC, Fujita MK, Schott RK, Thomas KN. 2021. Eye-body allometry across biphasic ontogeny in frog amphibians. Evol Ecol. 35(2):337–359. doi:10.1007/s10682-021-10102-3.

Stiebel-Kalish H, Reich E, Rainy N, Vatine G, Nisgav Y, Tovar A, Gothilf Y, Bach M. 2012. Gucy2f zebrafish knockdown – a model for Gucy2d-related leber congenital amaurosis. Eur J Hum Genet. 20(8):884–889. doi:10.1038/ejhg.2012.10.

Tamura K, Stecher G, Peterson D, Filipski A, Kumar S. 2013. MEGA6: Molecular Evolutionary Genetics Analysis Version 6.0. Molecular Biology and Evolution. 30(12):2725–2729. doi:10.1093/molbev/mst197.

Thomas KN, Gower DJ, Bell RC, Fujita MK, Schott RK, Streicher JW. 2020. Eye size and investment in frogs and toads correlate with adult habitat, activity pattern and breeding ecology. Proc R Soc B. 287(1935):20201393. doi:10.1098/rspb.2020.1393.

Thomas KN, Gower DJ, Streicher JW, Bell RC, Fujita MK, Schott RK, Liedtke HC, Haddad CFB, Becker CG, Cox CL, et al. 2022. Ecology drives patterns of spectral transmission in the ocular lenses of frogs and salamanders. Functional Ecology. 36(4):850–864. doi:10.1111/1365-2435.14018.

Van Nynatten A, Bloom D, Chang BSW, Lovejoy NR. 2015. Out of the blue: adaptive visual pigment evolution accompanies Amazon invasion. Biol Lett. 11(7):20150349. doi:10.1098/rsbl.2015.0349.

Van Nynatten A, Castiglione GM, De A. Gutierrez E, Lovejoy NR, Chang BSW. 2021. Recreated Ancestral Opsin Associated with Marine to Freshwater Croaker Invasion Reveals Kinetic and Spectral Adaptation. Molecular Biology and Evolution. 38(5):2076–2087. doi:10.1093/molbev/msab008.

Van Nynatten A, Duncan AT, Lauzon R, Sheldon TA, Chen SK, Lovejoy NR, Mandrak NE, Chang BSW. 2024. Adaptive Evolution of Nearctic Deepwater Fish Vision: Implications for Assessing Functional Variation for Conservation. Crandall K, editor. Molecular Biology and Evolution. 41(2):msae024. doi:10.1093/molbev/msae024.

Wake DB, Koo MS. 2018. Amphibians. Current Biology. 28(21):R1237–R1241. doi:10.1016/j.cub.2018.09.028.

Wan YC et al. 2023. Selection on Visual Opsin Genes in Diurnal Neotropical Frogs and Loss of the SWS2 Opsin in Poison Frogs. Molecular Biology and Evolution. 40(10):1–19

Weiss ER, Ducceschi MH, Horner TJ, Li A, Craft CM, Osawa S. 2001. Species-Specific Differences in Expression of G-Protein-Coupled Receptor Kinase GRK7 and GRK1 in Mammalian Cone Photoreceptor Cells: Implications for Cone Cell Phototransduction. J Neurosci. 21(23):9175–9184. doi:10.1523/JNEUROSCI.21-23-09175.2001.

Wertheim JO, Murrell B, Smith MD, Kosakovsky Pond SL, Scheffler K. 2015. RELAX: Detecting Relaxed Selection in a Phylogenetic Framework. Molecular Biology and Evolution. 32(3):820–832. doi:10.1093/molbev/msu400.

White ND, Batz ZA, Braun EL, Braun MJ, Carleton KL, Kimball RT, Swaroop A. 2022. A novel exome probe set captures phototransduction genes across birds (Aves) enabling efficient analysis of vision evolution. Molecular Ecology Resources. 22(2):587–601. doi:10.1111/1755-0998.13496.

Wisotsky SR, Kosakovsky Pond SL, Shank SD, Muse SV. 2020. Synonymous Site-to-Site Substitution Rate Variation Dramatically Inflates False Positive Rates of Selection Analyses: Ignore at Your Own Peril. Crandall K, editor. Molecular Biology and Evolution. 37(8):2430–2439. doi:10.1093/molbev/msaa037.

Wu Y, Hadly EA, Teng W, Hao Y, Liang W, Liu Y, Wang H. 2016. Retinal transcriptome sequencing sheds light on the adaptation to nocturnal and diurnal lifestyles in raptors. Sci Rep. 6(1):33578. doi:10.1038/srep33578.

Wu Y, Wang H, Hadly EA. 2017. Invasion of Ancestral Mammals into Dim-light Environments Inferred from Adaptive Evolution of the Phototransduction Genes. Sci Rep. 7(1):46542. doi:10.1038/srep46542.

Yang R-B, Robinson SW, Xiong W-H, Yau K-W, Birch DG, Garbers DL. 1999. Disruption of a Retinal Guanylyl Cyclase Gene Leads to Cone-Specific Dystrophy and Paradoxical Rod Behavior. J Neurosci. 19(14):5889–5897. doi:10.1523/JNEUROSCI.19-14-05889.1999.

Yang Z, Wong WSW, Nielsen R. 2005. Bayes Empirical Bayes Inference of Amino Acid Sites Under Positive Selection. Molecular Biology and Evolution. 22(4):1107–1118. doi:10.1093/molbev/msi097.

Yang Z. 2007. PAML 4: Phylogenetic Analysis by Maximum Likelihood. Molecular Biology and Evolution. 24(8):1586–1591. doi:10.1093/molbev/msm088.

Yohe LR, Abubakar R, Giordano C, Dumont E, Sears KE, Rossiter SJ, Dávalos LM. 2017. *Trpc2* pseudogenization dynamics in bats reveal ancestral vomeronasal signaling, then pervasive loss. Evolution. 71(4):923–935. doi:10.1111/evo.13187.

Yovanovich CAM, Grant T, Kelber A. 2019 Jan 1. Differences in ocular media transmittance among classical frog model species and its impact on visual sensitivity. Journal of Experimental Biology.:jeb.204271. doi:10.1242/jeb.204271.

Yovanovich CAM, Pierotti MER, Kelber A, Jorgewich-Cohen G, Ibáñez R, Grant T. 2020. Lens transmittance shapes ultraviolet sensitivity in the eyes of frogs from diverse ecological and phylogenetic backgrounds. Proc R Soc B. 287(1918):20192253. doi:10.1098/rspb.2019.2253.

