## Supplementary figures and images for "Evolutionary and ontogenetic shifts from aquatic to terrestrial light environments in frogs provide new insights into the vertebrate phototransduction cascade"

### Fig. S1

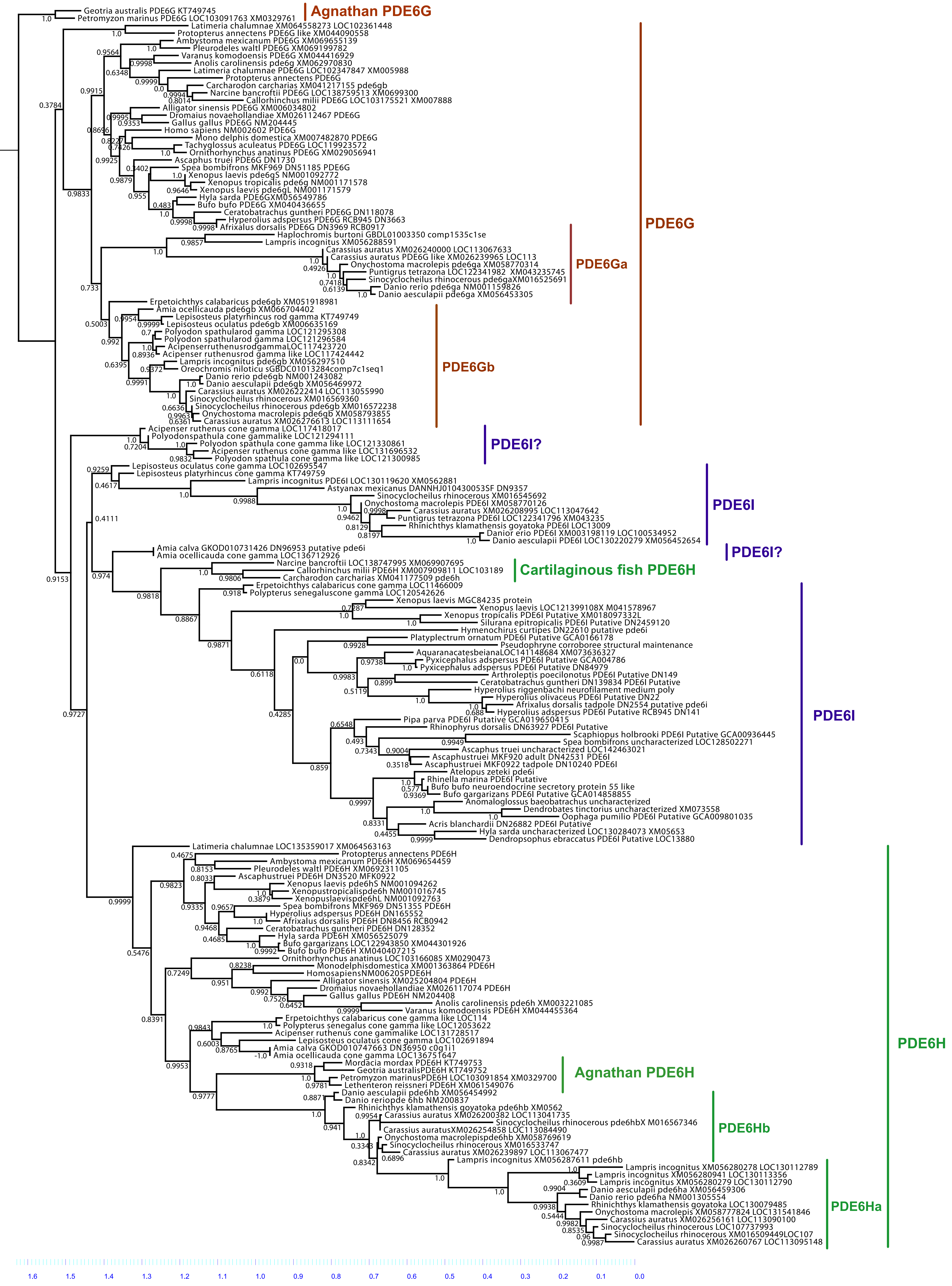


**Fig. S1. Maximum likelihood gene tree of PDE6 gama genes.**
